# Proposed network of Marine Protected Areas supports larval dispersal and connectivity in the Eastern Mediterranean

**DOI:** 10.1101/2024.04.28.591505

**Authors:** Igal Berenshtein, Nir Stern, Aviyam Tagar, Claire B. Paris, Omri Lapidot, Arseniy R. Morov, Erick Fredj, Jacob Zaken, Eli Biton

**Affiliations:** Department of Marine Biology, Leon H Charney School of Marine Sciences, University of Haifa, Haifa, Israel; Rosenstiel School of Marine, Atmospheric and Earth Sciences, University of Miami, Miami, USA; National Institute of Oceanography, Israel Oceanographic and Limnological Research (IOLR), Haifa, Israel; Department of Computer Science, Jerusalem Institute of Technology (JCT), Jerusalem, Israel

## Abstract

The marine environment of the Eastern Mediterranean is under growing threat due to natural and anthropogenic stressors. Networks of Marine Protected Areas (MPAs) are effective tools in protecting marine environments and conserving their biodiversity. Currently, only 4% of the Israeli territorial waters are declared as MPAs, however six new MPAs, which will encompass more than 20% of the Israeli territorial waters, are planned. A central component in the effectiveness of MPAs is the degree to which the protected populations are connected. The purpose of our study is performing a comprehensive connectivity analysis for the proposed network of MPAs. We find that the proposed network substantially supports local and regional larval connectivity patterns for five target species in terms of the number of recruits, betweenness centrality, as well as the number of regional and local MPAs connections. Overall, the results provide strong support for the efficiency of the proposed MPAs in facilitating local and regional larval connectivity. Our findings will be useful for marine spatial planning and natural resource management and will enhance the protection and conservation of our marine environment.

## Introduction

The Mediterranean marine ecosystem provides critically important ecosystem services from environmental, economic, and social aspects (Liquete *et al*., 2016). Yet at the same time, this system is under high anthropogenic pressures including fishing (FAO, 2016), pollution (Dong *et al*., 2022), invasive species (Galil *et al*., 2021) and climate change (Albano *et al*., 2021). As a result, marine natural resources in the Eastern Mediterranean have been substantially declining in the past decades (Demirel *et al*., 2020).

The Israeli territorial waters in the Mediterranean which cover more than 3,800 km^2^, harbor a diverse fauna, including multiple species of fish, invertebrates, sea turtles, marine mammals, and more. The sea is a crucially important asset from an environmental, economic, and social aspects, providing income, revenue, nutrition, recreation, clean air, and a viable source of drinking water. At the same time, this marine environment is under an increasing anthropogenic exploitation including over-fishing, natural gas infrastructure, desalination plants, and more. This pressure, together with global climate changes pose a severe threat to the existence and stability of this important asset, which warrants timely and appropriate protection(Ryabinin *et al*., 2019).

For effective protection of marine populations, MPAs must be: (1) self-sustaining, i.e. enable sufficient return of settlement-stage larvae to their natal MPA; or (2) sufficiently connected to other MPAs via dispersal; (3) replenish fishing areas outside the reserve *via* spill-over (Planes *et al*., 2009; Pelc *et al*., 2010). The connectivity pattern within the proposed MPAs in the Israeli Mediterranean is currently unknown. The common practice for a comprehensive examination of demographic connectivity is the use of biophysical models. These are typically individual-based Lagrangian frameworks combining oceanic currents data with species-specific biological traits, producing virtual larval trajectories used to compute population connectivity(Werner *et al*., 2007). In this paper we present a three-pronged biophysical approach focusing on the physical, biological, and biophysical components, computing an analysis of marine population connectivity pattern in the Eastern Mediterranean.

The circulation of the Israeli continental shelf and slope is a part of the general circulation of the Levantine basin, composed of the bifurcating Atlantic Ionian Stream, the Mid-Mediterranean Jet, and the cyclonic basin-wide current that follows the coast. During most of the year, the surface circulation is dominated by a cyclonic along-slope current flowing over the shelf and slope (Rosentraub and Brenner, 2007), though it may be interrupted by episodes of southward flow due to strong easterly winds and anticyclonic eddies off the shelf (Rosentraub and Brenner, 2007). Interactions between the shelf and shallow coastal areas result in complex circulation patterns over the shelf, e.g. the meandering of the along-slope current (Gertman *et al*., 2010), and the formations of eddies, and filaments of coastal waters intruding to deep waters (Efrati *et al*., 2013). The latter was demonstrated by the detachment of the anticyclonic Shikmona eddy near Haifa bay (Gertman *et al*., 2010). Over the slope, a seasonally varying along-slope baroclinic jet also appears during the summer and early winter. Beyond the slope area, the flow is characterized by a series dynamic meso-scale eddies (Amitai *et al*., 2010). These oceanic features play a key role in the transport of water masses along with their associated physical, chemical, and biological properties. One of these important biological properties include the transport of early life stages of marine organisms, or propagules (Cowen and Sponaugle, 2009).

Most benthic marine organisms are characterized by a bipartite life cycle: the adult stage, which is associated with a benthic habitat, and the larval stage, during which larvae set out to the open sea for a few days up to a few weeks, at the end of which they metamorphose and settle in their habitat (Gaines and Lafferty, 1995). The larval stage is the main dispersal mechanism of these organisms, and hence is of great importance as it determines population connectivity and meta-population dynamics(Cowen *et al*., 2000); consequently, affecting marine resources management strategies and MPAs network design (Palumbi, 2001).

The proposed network of MPAs is designed to protect representative and unique habitats in the Israeli Mediterranean marine environment: soft-sand, rocky, silt and clay, canyons, underwater eolianite cliffs; as well as other unique benthic marine habitats (Yahel and Engert, 2012). The different habitats that are characterized by different key species, which produce species-specific dispersal patterns according to their larval traits (Holstein *et al*., 2014; Wolanski and Kingsford, 2014; Berenshtein *et al*., 2016).

There is little available information regarding the regional MPA connectivity. (Andrello *et al*., 2013) demonstrated low connectivity for *Epinephelus marginatus* between MPAs in the entire Mediterranean, but relatively high connectivity within the Eastern Mediterranean MPAs. Notably, Eastern Mediterranean MPAs supplied larvae to the downstream Aegean sea, but not vice versa (Andrello *et al*., 2013). Yet, high resolution, regional scale comprehensive connectivity analysis which includes the proposed MPAs is missing.

Larval recruitment and connectivity are primary drivers governing marine population and stock dynamics (Hjort, 1914; Houde, 2008), and as such are increasingly being considered in the study and conservation of marine ecosystems, and in particular in the design of MPAs networks due to the importance of location (Kough *et al*., 2018; Balbar and Metaxas, 2019). In the current paper we performed a comprehensive connectivity analysis for the Eastern Mediterranean, focusing on the proposed Israeli MPAs network. Specifically, we have chosen five focal species, and examined the contribution of MPAs network to their regional and local connectivity patterns, computing connectivity-related estimates, such as number of connections and betweenness centrality. In addition, we performed a sensitivity analysis, covering realistic ranges of larval traits, such as larval swimming speeds and mortality coefficient estimates.

## Methods

Our methodology is based on a three-pronged approach: (1) Physical oceanography, which includes a nested ocean model simulating four years of data (2017-2020), validated with in-situ observations. (2) Biological component, which includes ichthyoplankton sampling, as well as a literature review, choosing target species, and providing the best available data regarding these species in the MPAs and their biologically relevant traits. (3) Bio-physical simulations which combines the biological and the physical components producing 3D larval trajectories. These trajectories were then used to analyze the connectivity patterns of the proposed MPAs. Given the physical and biological variation in the system, a sensitivity analysis was performed, covering the meaningful ranges of parameter values representing the spectrum of realistic connectivity patterns. In addition, we included a user-friendly interface, which displays the respective plots and results.

The physical component includes a nested model, with Copernicus Marine Environment Monitoring Service (CMEMS) as the outer model and South Eastern Levantine Israeli Prediction System (SELIPS) as the inner model. The CMEMS currents are based on hydrodynamic model currents output implemented over the Mediterranean Basin with a horizontal grid resolution of 1/24’ (∼ 4 km) and have 141 vertical levels (Figure. 1). The hydrodynamics are supplied by the Nucleus for European Modelling of the Ocean (NEMO; Clementi et al., 2016). The model includes data assimilation schemes of temperature and salinity vertical profiles, as well as those of satellite Sea Level Anomaly observations.

SELIPS is a sub-regional high resolution circulation model operated by Israel Oceanographic and Limnological Research (IOLR) that generates daily forecasts of temperature, salinity, and sea currents in the southeastern region of the Levantine basin (Figure 1). The oceanic general model used to run SELIPS is the Princeton Ocean Model (POM) ; Blumberg and Mellor, 1987). The resolution is 0.01°x0.00833° (about 1 km) with 27 sigma levels, and the minimal depth is 5 meters. Initial and boundary conditions of temperature, salinity, and water velocity are taken from ALERMO forecast system (Zafirakou-Koulouris *et al*., 2012).

Atmospheric fluxes of heat, fresh water, and momentum at the sea surface are computed from bulk formulas using the conditions taken from SKIRON, a regional atmospheric model forecast, including air temperature, specific humidity, air pressure, wind, total cloud cover and precipitation. Modeled heat flux was corrected using the remotely sensed sea surface temperature (SST) fields taken from Copernicus Marine Service. The validation of SELIPS model was carried out against in-situ measurements of temperature, salinity, and currents direction and velocity from Ashkelon, Hadera, and DEEPLEV mooring (for more details see Supplementary section S1).

**Figure 1.**
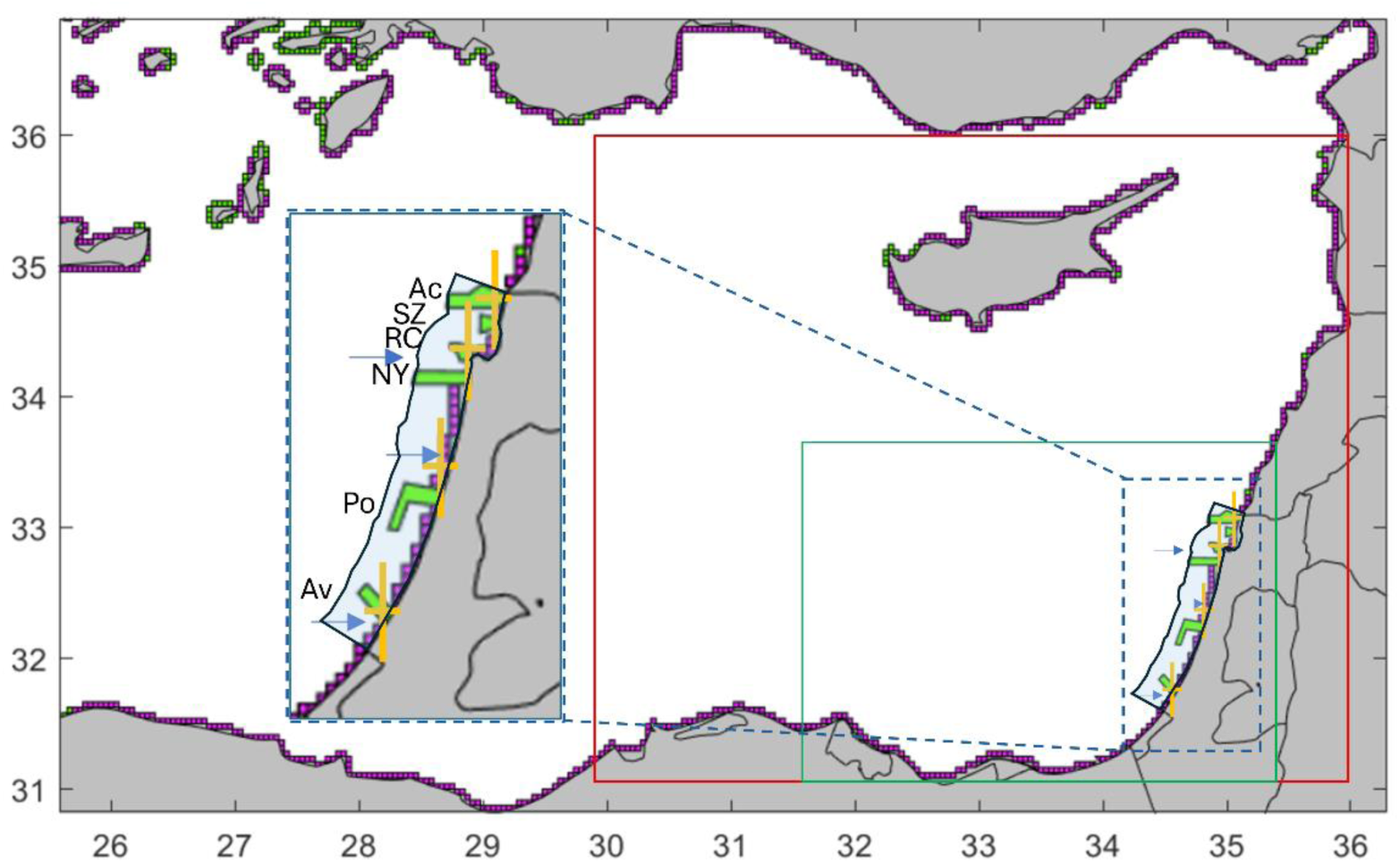
The modeled domain indicates the nested ocean models, coastal polygons, MPAs, and biological and hydrographic sampling stations. The outer ocean model is Copernicus Marine Environment Monitoring Service (CMEMS; red), and the inner ocean model is South Eastern Levantine Israeli Prediction System (SELIPS; green). Purple and green polygons represent regular (continental shelf) and MPAs regions along the coast. Cyan shaded area represents the Israeli territorial waters. Orange crosses represent the biological sampling sites from north to south: Achziv, Shikmona, Gdor, and Avtach. Green polygons represent proposed large MPAs of: Avtach (Av), Poleg (Po), Neve Yam (NY), Rosh Carmel (RC), Shavei Zion (SZ), and Achziv (Ac). Blue arrows represent the stations for hydrographic data measurements (temperature, salinity, current velocity, and strength). Note that for visualization clarity purposes, coastal polygons are small, however, in the simulations, coastal polygons were elongated to approx.. 20 km to cover the continental shelf.

The biological component included ichthyoplankton sampling scheme and a literature review. Ichthyoplankton samples were collected from four MPAs along the Eastern Mediterranean coast of Israel: Achziv, Shikmona, Gdor, and Avtach (Fig. 1). Ichthyoplankton sampling was conducted during spring to summer (May-August), except for the first sampling cruise of Achziv in august 2019 and the second sampling cruise of Michmoret in November 2020. Each sampling cruise was conducted during the night (from dusk to dawn). In total, sixteen cruises were conducted over the research period.

Target species were selected based on their ecological and economical importance, and a wide range of their life history traits is represented in our simulation scenarios (Table 1). In addition, the dispersal of some species may not be represented by the ichthyoplankton samples, as these may include exceptional larval dispersal dynamics (e.g., larvae from demersal spawning trapped in the bottom boundary layer (Bell and Brown, 1995). Hence selected species included those that have ecological and economical importance and appeared in the ichthyoplankton samples (except for *Scyllarides latus,* as samples were focused on ichthyoplankton and not invertebrate zooplankton). The following species were chosen for explicit species-specific representation in our simulations:

- *Epinephelus costae* (Goldblotch Grouper): a native species that can mainly be found on coastal rocky habitats, and feeds on mollusks and small fishes (Trophic Level (TL) = 3.9). It feeds on mollusks and small fishes (Trophic Level (TL) = 3.9). *E. costae* is mainly abundant in rocky habitats.
- *Nemipterus randalli* (Randal threadfin seabream): an invasive Red Sea Indo-Pacific species commercially important. *N. randalli* is a demersal benthic invertebrate feeder (TL = 3.8), highly abundant in sandy and muddy habitats along the continental shelf of the Levant Basin.
- *Parupeneus forsskali* (Red Sea goatfish): an invasive Red Sea species, with increasing commercial importance. *P. forsskali* is a benthic invertebrate feeder (TL = 3.5), abundant in shallow sandy and rocky habitats.
- *Mugil cephalus* (Flathead grey mullet): a commercially important native forage species, feeding on detritus, micro-algae and benthic organisms (TL = 2.5), and supporting higher trophic level predators. *M. cephalus* is abundant in shallow coastal regions, in sandy and rocky habitats.
- *Scyllarides latus* (Mediterranean slipper lobster)*: S. latus* is a native species, associated with rocky or sandy substrates at depths of 4–100 m. As this is an invertebrate species, there was no attempt to collect and identify this species in our ichthyoplankton sampling campaign. However, its importance was highlighted by managers, and was therefore included in the simulations.

### Biophysical component

Connectivity Modeling System (CMS, Paris et al., 2013) simulations using the nested model of CMEMS and SELIPS were run, performing general simulations representing five commercially and ecologically important species (Table 1), as well as general simulations, representing a realistic range of larval traits (Table 2). The model output was then further analyzed to compare between a situation with proposed MPAs vs. without MPAs, computing: recruitment success, self-recruitment, the number of recruits originated from Israelis waters (Israeli waters, herein ‘locally’), total number of recruits locally settled, the number of recruits originated and settled locally, the number of local connections, and the number of regional MPAs connections, number of recruits in MPAs (regional). In addition, we computed betweenness centrality, which measures the proportion of shortest paths between nodes that pass through a given node, which represents the degree by which MPAs act as gateways of dispersal (Andrello *et al*., 2013).

The efficiency of MPAs was based on previously published empirical evidence for increased reproductive output, i.e., number of produced larvae (Marshall *et al*., 2019) and higher biomass (Sala and Giakoumi, 2018) in MPAs compared to unprotected areas. For scenarios which considered effective MPAs, we used an estimate of seven-fold more reproductive output (i.e., number of released larvae) in MPAs compared to non-protected regions (Marshall et al., 2019, Sala and Giakoumi, 2018). Note that the reproductive output is proportionate to the area of the habitat (or polygon). The control scenarios against which the formers were compared to non-protected areas in terms of reproductive output. The depth of released virtual larvae were both near the surface (depth=5 m) and in the water column, chosen as the mid-depth between the bottom and the water surface per release location. Life history traits that were used for model parameterization include ontogenetic swimming speeds, pelagic larval duration (PLD), mortality rates (K), and shoreward swimming orientation modified through the concentration parameter (κ) of swimming directions around the shoreward direction (Table 2).

**Table 1.** Life history traits for the biophysical model based on the nested model of CMEMS and SELIP for four years: 2017-2020. Spawning season is simulated as May-September in accordance to the species estimated spawning season (Stern, 2016).

| *Species* | Common name | Origin | Habitat | Pelagic Larval Duration (PLD, days) | References |
| --- | --- | --- | --- | --- | --- |
| *Nemipterus randalli* | Randal threadfin seabream | Non-indigenous | Sand | 21 | (Hall *et al.*, 2019) |
| *Parupeneus forsskali* | Red Sea goatfish | Non-indigenous | Sand / Shallow rocky | 25-37 | (Leis and McCormick, 2002) |
| *Mugil cephalus* | Flathead grey mullet | Native | Shallow / Rocky / Sand | 42 | (Ditty and Shaw, 1996) |
| *Epinephelus costae* | Goldblotch Grouper | Native | Rocky | 20-40 | (Andrello *et al.*, 2013) |
| *Scyllarides latus* | Mediterranean slipper lobster | Native | Rocky | 200-300 | (Martins, 1985) |

**Table 2.** Scenarios of CMS biophysical run for computing general connectivity patterns based on the nested model of CMEMS and SELIP for four years: 2017-2020. Parameters include Pelagic Larval Duration (PLD), in days’ units, instantaneous mortality rate, relative egg production, and swimming speed (cm/s). Kappa represents the concentration parameter of the von-Mises distribution from which virtual larval swimming directions are drawn from.

| Parameter | low | medium | high | references |
| --- | --- | --- | --- | --- |
| PLD (d) | 2-5 | 21-40 | 200-300 | (Shanks, 2009) |
| Mortality rate | 0 | 0.2 | 0.4 | (Houde, 1989) |
| Egg production level | x1 | NA | x7 | (Marshall *et al.*, 2019) |
| Swimming speed (cm/s) | 0 | 15 | 30 | (Fisher and Bellwood, 2003; Fisher, 2005; Fisher and Leis, 2010) |
| Orientation ($\kappa$) | 0 | 2.5 | 5 | (Codling *et al.*, 2004; Staatterman *et al.*, 2012; Berenshtein *et al.*, 2016, 2018b) |

## Results

From a regional perspective, the results for larval fish with average PLD (e.g., *Parupeneus forsskali*) demonstrate high connectivity with no apparent dispersal barriers (Fig. 2A). In addition, an a-symmetric localized recruitment is evident, corresponding to the general counter-clockwise circulation of the Eastern Mediterranean (Rosentraub and Brenner, 2007). Specifically, Egyptian coastline supplies larval output mainly to itself, but also to Israeli waters; Israel coastal habitats supplies virtual larvae mainly to itself but also to Lebanon coastal habitats; Lebanon supplies to itself but also to Syria, and so on and so forth following a general “anti-clockwise” direction of supply, corresponding to the regional general circulation pattern (Rosentraub and Brenner, 2007). The degree of connectivity and length of dispersal increases with PLD, such that short PLDs (1-2 d), the most dominant feature is the diagonal in the connectivity matrix (Figure 2C), and the connections are mainly local (Fig. 2D-F). For moderate PLDs (e.g., *P. forsskali* ; PLD= 25-37 d; Fig. 2A-B), there is a higher degree of spread of connections (Fig. 2A-B), and for long PLDs (e.g., *Scyllarides latus*; PLD= 200-300 d; Fig. 2H-J), the connections further increase, connecting between much farther locations, and reducing the degree of self-recruitment, obstructing the diagonal feature in the connectivity matrices (Fig. 2H-J).

**Figure 2.**
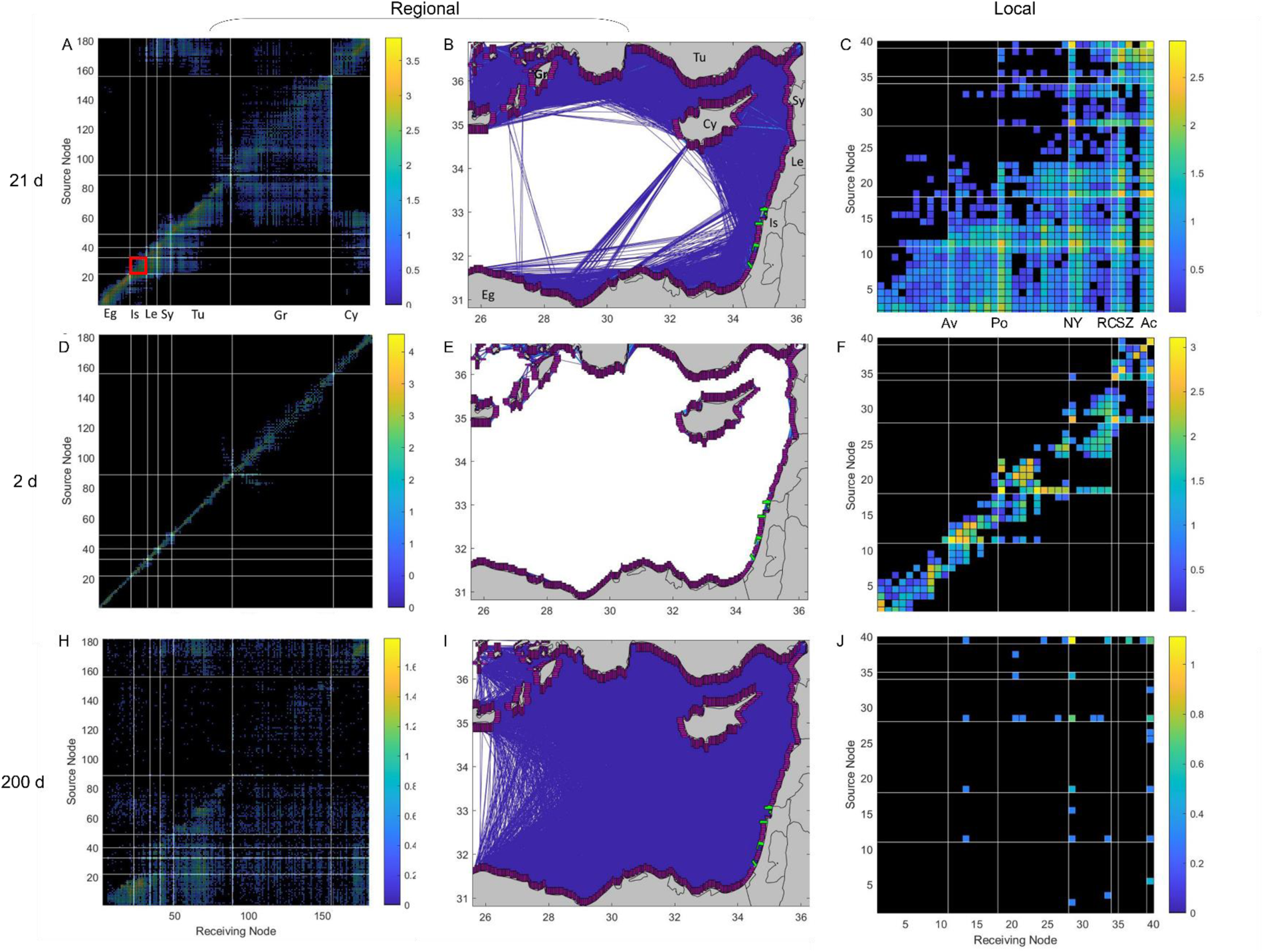
Regional (left and middle panels) and local (right panel) connectivity results for species with variable Pelagic Larval Durations (PLD) of 21-40 d (top row), PLD= 1-2 d (middle row) and, PLD= 200-300 d (bottom row), simulations were run for the years 2017-2020. Regional connectivity matrices, in which area between two adjacent white lines (both vertical and horizontal) represent the coastal/ continental shelf areas. Numbers in the left panels’ connectivity matrices represent continental shelf polygons, counted from the southwestern area of the domain, i.e., west Egypt (Eg), counterclockwise through: Israel (Is), Lebanon (Le), Syria (Sy), Turkey (Tu), Greece (Gr), and Cyprus (Cy). White lines in the right panels (local-Israel), represent the proposed large MPAs of Avtach (Av), Poleg (Po), Neve Yam (NY), Rosh Carmel (RC), Shavei Zion (SZ), and Achziv (Ac), ordered from bottom to top and from left to right. Colorbars represent log_10_ transformation to virtual larval numbers (for more details about the simulation see Table 3). Scenarios here represent baseline conditions in which MPAs do not contribute more virtual larvae (per area) than other coastal/continental shelf regions. Red square in panel A represents the local (Israeli) connectivity matrix presented in panels C, F, and J.

The main larval sinks for the cumulative connectivity matrix (summed over all scenarios from Table 3) of the proposed MPAs is mainly in Israel, with contributions to Lebanon, Syria, Turkey, and Greece (Fig. 3A,C). The main larval sources to the proposed MPAs originate from Israel and Egypt (Fig. 3B,D). Notably, the proposed MPAs (Fig. 3C) contribute more virtual larvae to Lebanon than to Israel, mainly because of the contribution of the northern MPAs (Achziv, Poleg, and Nve-Yam). The southern MPAs contribute more larvae to the Israeli region.

**Figure 3.**
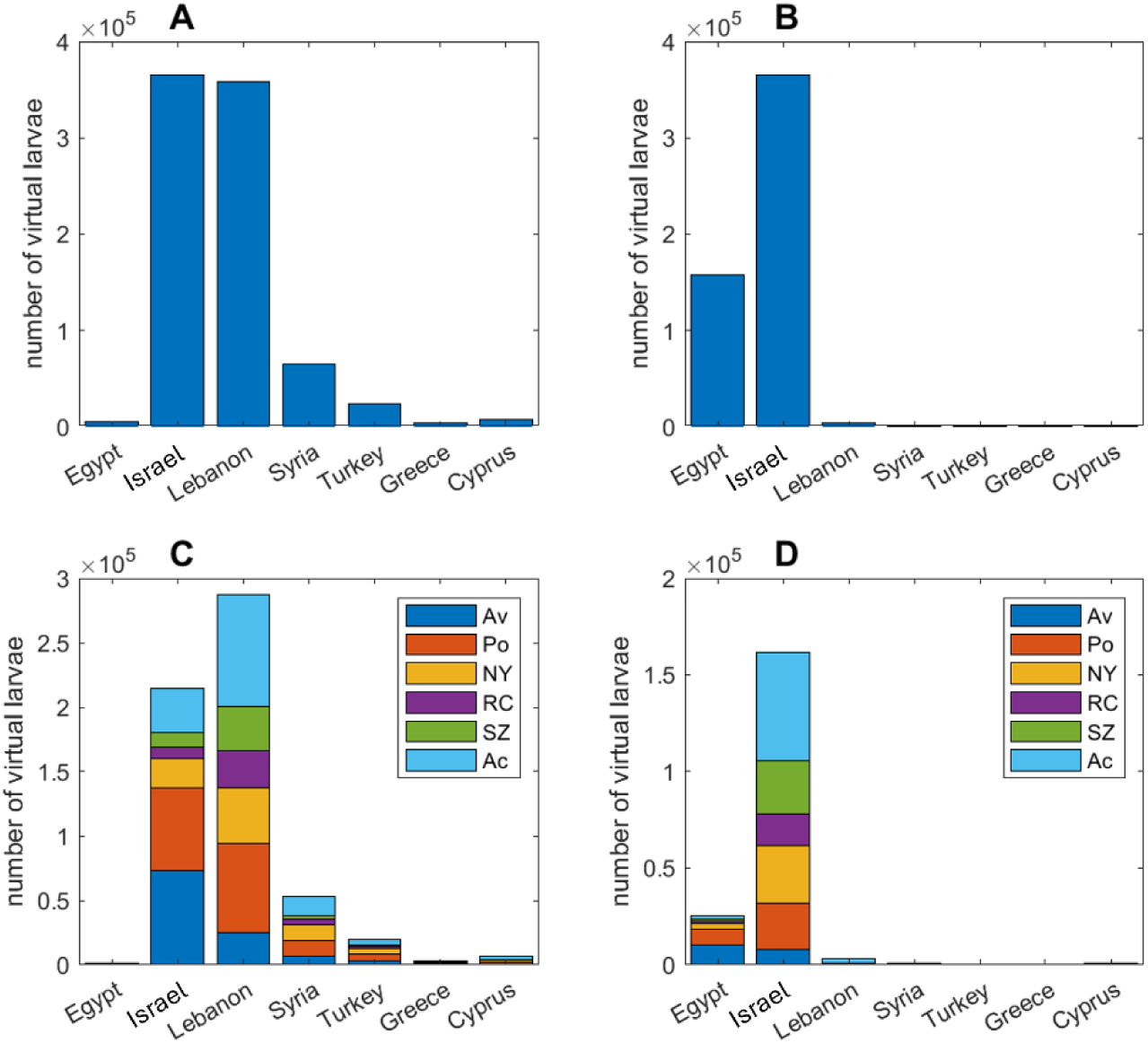
The sinks (A,C) and sources (B,D) with respect to Israel as a whole (upper panels) and the proposed MPAs (bottom panels) for the cumulative connectivity matrix summed across all the simulations scenarios (Table C1,2). Countries included are Egypt (Eg), Israel (Is), Lebanon (Le), Syria (Sy), Turkey (Tu), Greece (Gr), and Cyprus (Cy); suggested large MPAs are: Avtach (Av), Poleg (Po), Neve Yam (NY), Rosh Carmel (RC), Shavei Zion (SZ), and Achziv (Ac).

We find that effective MPAs substantially increase connectivity related estimates of recruitment success (+9.5±31%), recruitment success in MPAs (+335±238%), number of connections (+21±49%), regional MPA connections (+70±106%), local MPA connections (+35±78%), no. of recruits in Israel originated from Israel (218±238%), and betweenness centrality (+76±112%) (Table 3). The effect of MPAs on regional MPA connectivity is evident both in the strength and the number of connections (Fig. 4).

**Figure 4.**
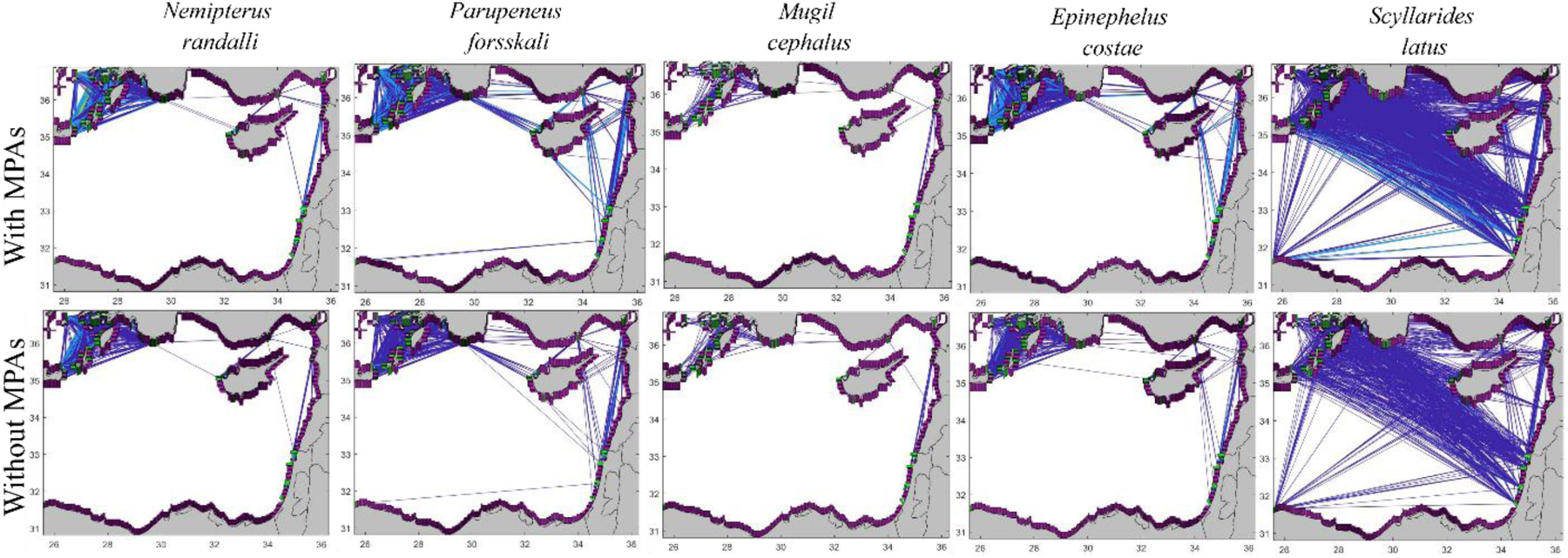
Regional contribution of MPAs. Upper and lower panels represent simulations in MPAs are considered as regular habitat polygons in terms of larval production (upper panels), and effective MPAs with increased larval production (x7) (lower panels). Lines represent connections between MPAs in the eastern Mediterranean with warmer line colors represent more connections between the MPAs.

The contribution of the proposed MPAs varied between the different connectivity estimates. The MPAs with the highest contribution in terms of betweenness centrality estimates from a local perspective were Poleg and Neve Yam (also known as Yam Karmel) (Fig. 5A), and from a regional perspective: Avtach, Poleg and Neve Yam (Fig. 5C). Achziv is the MPAs which provides settlement habitat for most larvae both from local and regional perspectives (Fig. 5B,E). Avtach is the MPA that produces the highest number of larvae which recruit both locally and regionally (Fig. 5C,F), with Achziv providing the second highest contribution of larvae from a regional perspective (Fig. 5F).

**Figure 5.**
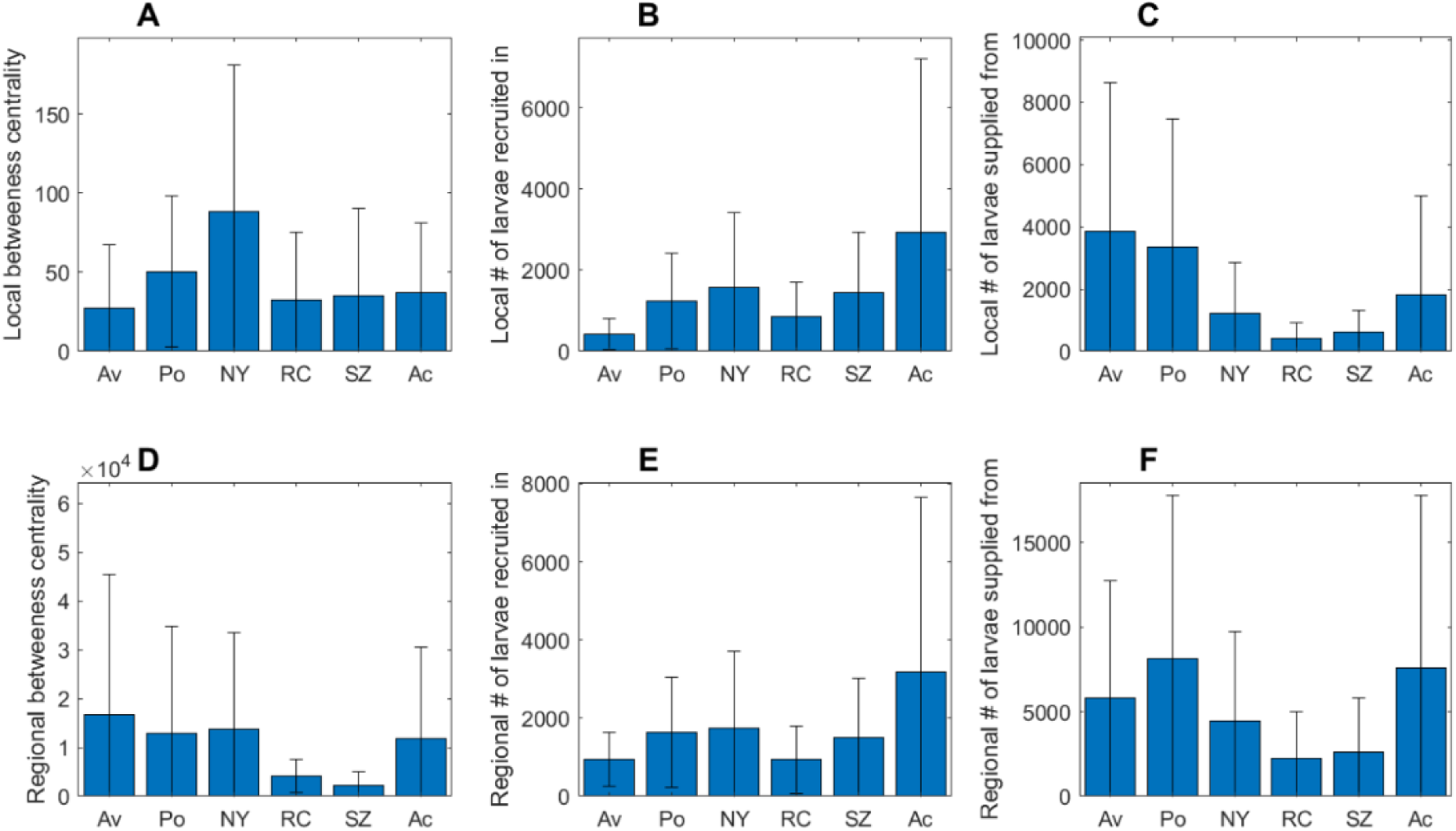
The contribution of individual MPAs to local (upper panels) and regional (lower panels) connectivity estimates: (A,C) betweenness centrality, (B,E) number of virtual larvae recruited in the MPA, and (C,D) number of virtual larvae from the MPA recruited elsewhere. Suggested large MPAs: Avtach (Av), Poleg (Po), Neve Yam (NY), Rosh Carmel (RC), Shavei Zion (SZ), and Achziv (Ac). Local-considers the Israeli region only, and Regional – considers the entire Eastern Mediterranean domain.

Our results demonstrate that increase in orientation parameter (*κ= 5*) and swimming speeds (terminal speed = 30 cm/s) substantially increase recruitment success (+115%), recruitment success in MPAs (+62%), number of connections (+33%), and number of connections in MPAs (+38%) (Table 3), as was shown in previous studies (Staaterman *et al*., 2012; Berenshtein *et al*., 2018a, 2018b). An opposite trend is evident with the implementation of larval mortality (K=0.2), resulting in drastic declines in connectivity estimates of recruitment success in MPAs (-97%), number of connections (-87%), number of connections in MPAs (-85%) (Table 3), as was shown previously for other systems (Berenshtein *et al*., 2018a).

**Table 3.** Simulation scenarios and their results. Scenarios names with MPAs represent scenarios in which larger number (x7) of eggs/larvae are released in MPAs. Columns titles abbreviations: recruitment success (recSuc), MPAs recruitment success (recSuc_MPA), number of connections (Conn_num), number of connections between MPAs (Conn_num_MPA), number of connections in Israel (Conn_num_Isr), number of recruits that originated from and settled in Israeli coastal habitats (Num_of_rec_in_from_Isr), Betweenness centrality (BetCen), mean number of recruits that originated from and settled in coastal habitats (SelRec).

| Scenario | recSuc | recSuc_MPA | Conn_num | Conn_num_MPA | Conn_num_Isr | Num_of_rec_in_from_Isr | BetCen | SelRec |
| --- | --- | --- | --- | --- | --- | --- | --- | --- |
| *Nemipterus randalli* | 0.18 | 44516 | 65051 | 3493 | 951 | 26591 | 42.81 | 0.074 |
| *Parupeneus forsskali* | 0.15 | 34479 | 84419 | 3720 | 956 | 22390 | 24.99 | 0.050 |
| *Mugil cephalus* | 0.12 | 26439 | 34359 | 3190 | 755 | 12582 | 52.64 | 0.090 |
| *Epinephelus costae* | 0.01 | 865 | 14230 | 326 | 594 | 2840 | 17.31 | 0.016 |
| *Scyllarides latus* | 0.02 | 723 | 18202 | 570 | 33 | 58 | 23.83 | 0.006 |
| *Nemipterus randalli* (MPAs) | 0.14 | 1.54E+05 | 51499 | 3604 | 847 | 50794 | 74.63 | 0.072 |
| *Parupeneus forsskali* (MPAs) | 0.13 | 1.35E+05 | 72038 | 4809 | 872 | 37056 | 62.79 | 0.047 |
| *Mugil cephalus* (MPAs) | 0.16 | 1.85E+05 | 67290 | 4812 | 923 | 59171 | 18.99 | 0.066 |
| *Epinephelus costae* (MPAs) | 0.01 | 917 | 14187 | 356 | 592 | 2873 | 17.67 | 0.015 |
| *Scyllarides latus* (MPAs) | 0.03 | 4566 | 26238 | 2032 | 90 | 384 | 75.11 | 0.008 |
| 2_0_0_0 | 0.06 | 57379 | 6713 | 749 | 238 | 18200 | 65.64 | 0.223 |
| 20_0_0_0 | 0.07 | 23953 | 41202 | 2656 | 828 | 15913 | 63.75 | 0.043 |
| 20_0_2.5_30 | 0.15 | 38693 | 54659 | 3599 | 1116 | 26837 | 38.7 | 0.024 |
| 20_0_5_15 | 0.12 | 32924 | 48674 | 3373 | 1073 | 25334 | 30.32 | 0.030 |
| 20_0_5_30 | 0.15 | 38695 | 54859 | 3649 | 1120 | 26801 | 17.53 | 0.024 |
| 20_0.2_0_0 | 0 | 1042 | 6822 | 551 | 340 | 738 | 32.63 | 0.049 |
| 20_0.4_0_0 | 0 | 49 | 413 | 46 | 30 | 33 | 4 | 0.071 |
| 40_0_0_0 | 0.07 | 17668 | 58357 | 2856 | 794 | 9999 | 121.85 | 0.029 |

## Discussion

In this research we formulated a three-pronged framework for effectively quantifying the local and regional connectivity patterns, focusing on the potential contribution of proposed MPAs to these patterns. The physical and biological components formed the essential inputs based on which the biophysical/synthesis component computed larval trajectories and connectivity. Our results demonstrate high regional and local connectivity, and strongly support the effectiveness of the proposed MPAs from local and regional perspectives. This trend was consistent across the entire range of realistic larval traits, except for very short PLDs (2 d), and application of high mortality coefficient (Table 3). The high connectivity estimated between the different continental shelf polygons in Israel across a variable range of PLDs (Fig. 2), and the high level of Israeli-local self-subsidy (21%; Figs. 2, 3) suggests that protection of marine areas will be beneficial and effective in supporting local marine ecosystems. The contribution of the MPAs to regional MPA connectivity (Figs. 3, 4) and regional betweenness centrality (Fig. 5), highlights the regional importance of the protected areas.

The proposed MPAs network seems to be effective for short PLD scenarios (PLD=2 d; Fig. 2D-F), such that all proposed MPAs provided larvae to at least one adjacent MPA (Fig. 2F). In addition, the distance between the MPAs represents an optimal “rule-of-thumb” distance (20-200 km) according to previous analyses (Kinlan and Gaines, 2003; Halpern *et al*., 2006). The proposed design of large MPAs supports both short dispersal marine organisms through self-subsidy, as well as through adjacent MPAs which, especially in the northern area, are closely spaced (∼10-15 km nearest-neighbor distance between Neve Yam, Rosh Carmel, Shavei Zion, and Achziv). Because the proposed MPAs are large, long (∼20 km) and wide (>6 km), and cover the entire depths of the territorial waters, they provide a basis for self-settlement for species with short PLDs, as well as large area for production and settlement of larvae. The coverage of the depth gradient allows the inclusion of a variety of coastal and deep-sea habitats and enables their connectivity. In addition, the width of the MPAs alleviates the “edge effect”, which reduces the benefits of MPAs (Ohayon *et al*., 2021).

Our results indicate high local and regional connectivity, regardless of the MPAs. However, the degree of connectivity in the field is dependent on the abundance and diversity of the larvae/eggs released. Indeed, our ichthyoplankton sampling demonstrated the occurrence of eggs and settlement-stage larvae in all sampled sites indicates that these sites act both as sources (eggs) and settlement destination for larval fish (Fig. B1,2) as is simulated in our scenarios. In addition, the fact that species composition was mixed among sites (Fig. B5) supports our estimations of high connectivity. Hence, the protection of vast regions from fishing and/or other harmful activities will also enhance the degree of connectivity. This is even more substantial when considering the effect of the non-linear relationship between size and fecundity, also termed as BOFFFFs (Big, old, fat, fertile female fish; (Hixon et al., 2014)). Protection from fishing enables the growth of such mega-producers, which boosts the total production of larvae and connectivity.

However, this apparent advantage, when viewed in the spectacles of natural marine ecosystem, may become a “two-edged sword” when considering the connectivity of invasive species or the propagation of marine pollutants across our domain. Indeed, the “successful invasion” of Lessepsian migrates with pelagic propagules in the Eastern Mediterranean agrees with our “high regional and local connectivity” results (Lasram *et al*., 2008). Similarly, the tar pollution which occurred in the Israeli shores during February 2021, demonstrated that a slick with limited dimensions (narrow oil slick strip of 67x3 km) identified 50 km west of Ashkelon, dispersed across the entire Israeli shores within a week. Nevertheless, it has been shown repeatedly that effective MPAs contribute to the resilience of marine ecosystems, which in turn, increases their capacity to overcome disturbances such as bioinvasions and marine pollution events (Kimbro *et al*., 2013; Mellin *et al*., 2016; Chagaris *et al*., 2020).

The three-pronged framework which we created as part of this research, together with additional evidence, provides a powerful tool for studying local and regional connectivity. However, this methodology has multiple limitations and consists of multiple levels of uncertainty. For example, the computation of the current fields is limited in its resolution in time and space and does not capture fine-scale and possibly small sub-mesoscale features. In addition, the parameterization of Eddie diffusivity in our model (as well as in nearly all larval dispersal models) is isotropic across space and time, which is non-realistic (North *et al*., 2009). Similarly, the boundary conditions of the model which are forced by regional atmospheric and oceanic models, themselves consist of uncertainty and limitations. Nevertheless, the ocean model agrees with the in-situ hydrographic measurements of temperature, salinity, and currents at multiple locations in the domain, across extensive periods of time (Supplementary section S1).

Uncertainties and limitations in the biological component mainly include limited ichthyoplankton sampling in time and space due to budget and personnel limitations. Specifically, while the MPAs extend ∼20 km offshore, our sampling effort was limited to a few kilometers offshore, hence the habitats associated with deep and open waters (maximal depth ∼900m) were not sampled. Nevertheless, more than 120 species were identified in the larval pool, among which a few were chosen as the target species for the simulations. Biological information that could be highly relevant for future studies is the locations and characteristics of groupers’ spawning aggregations, and the orientation behavior of estuary/coastal associated species such as *M. cephalus*.

The uncertainties in the biological and physical components are propagated to the biophysical model, which adds its own set of uncertainties, including the application of isotropic stochastic mortality, which likely differs across space and time, and implementation of a deterministic swimming speed-at-age equation, without accounting for individual variation. Recent advances in the understanding of larval orientation may enable a more accurate future implementation of larval orientation behavior (Berenshtein *et al*., 2022). Explicit Information about larval behavior may be critical because even slight changes in larval traits such as orientation parameter and swimming speeds, may result in drastic changes in larval dispersal outcome. Lastly, while our analyses indicate connectivity trends within a single generation, it is in fact an ongoing process across multiple generations and locations. A good future research avenue would be to compare our resulted connectivity patterns to genetic connectivity estimates, as was recently carried out in the central Mediterranean (Legrand *et al*., 2022). Additional information that could shed light on the materialized connectivity patters is the comparison of species composition, and number of common species as was applied elsewhere (Paris *et al*., 2020).

Marine ecosystems’ resilience is critically dependent on larval connectivity, and it is especially crucial during this period of rapid degradation of marine ecosystems and loss of biodiversity (Ryabinin *et al*., 2019). Given the importance of larval stages, and the fact that the local knowledge about their traits and sensitivity to environmental changes is scarce, more focus is needed on studying local and regional larval ecology. Similarly, the effect of multiple stressors, and the exploration of their effects *via* integrative modeling frameworks (Berenshtein *et al*., 2020) should be further studied and appropriate frameworks should be developed and implemented accordingly (Shropshire *et al*., 2022). As long as such major knowledge gaps exist, protecting large areas from fishing and other threats would remain the safest strategy to mitigate unknown sources of future connectivity and recruitment failures.

## Conclusions

We conclude that:

- The proposed network of MPAs is effective in supporting local and larval dispersal and connectivity.
- This would be effective under the condition that no-take zones will indeed be enforced, and disruptive anthropogenic activities will be prohibited.
- Overall, the proposed MPAs network can serve as our “insurance policy” or “larval bank” against anticipated and unanticipated future stressors.

## Acknowledgements

We would like to thank Prof. Gil Rilov for the valuable scientific advice with respect to this project, and the co-supervision of Aviyam Tagar M.Sc. degree. We would also like to thank Dr. Arseniy Morov for his technical support in the research cruises, molecular work, and other aspects of this project. We thank the University of Miami Institute for Data Science and Computing (IDSC) for the use of the Pegasus cluster.

## Supplementary Information

### Supplementary section S1 – SELIPS validation

In order to examine the performance of the SELIPS model, SELIPS results were compared with measurements in the coastal environment, and in the open sea. These included currents) temperature and salinity data obtained from the following sources: 1 Two coastal measurement stations located at the ends of the coal piers of the Orot Rabin and Rotenberg power plants in) Hadera and Ashkelon, which are about 2.2 km from the coastline and at a depth of 26 m; 2 The DEEPLEV) mooring) sampling station is located about 45 km northwest of the Haifa coast and at a depth of about 1600 m.

#### Comparison With stations in Hadera and Ashkelon

Overall, the SELIPS results and the temperature and salinity values measured at the measurement stations in Ashkelon (Lon 34.49917, Lat 31.63472) and Hadera (Lon 34.863070, Lat 32.470530) (Figures A1-4) are well matched as computed using root-mean-square error (RSME). The results of the model closely followed the seasonal dynamics of temperature and salinity. However, in the case of the salinity values, the observed seasonal amplitude is greater and indicates values higher in summer and lower in winter than those obtained based on the results of the model. A possible explanation for these discrepancies is related to factors that are not included. The model runs include the contribution of increased sedimentation of the streams during the winter, particularly during winter storms accompanied by heavy rainfall, and on the other hand, a possible contribution of brine flow from the desalination plants adjacent to the measuring stations. The comparisons between the streams measured in Ashkelon and Hadera and the results of the model are provided in Figures A5-8 and 6-10, respectively. In all cases, the flow observations were taken from a depth of 5 meters, and were taken every three hours to match the model’s output. Both the observations of the currents in Ashkelon and Hadera and the results of the model indicate that the coastal flow moves mainly along the coastline, with a clear predominance towards the north. Overall, the results of the model follow the data measured in Hadera nicely (the Fourier norm stands at 0.78), but there is a slight shift of about 10 degrees compared to the data, which apparently originated from the bias of the model and the direction of its coastline compared to the direction of isobaths adjacent to the measuring stations. At the Ashkelon station, it can also be seen that the model tracks seasonal changes in the intensity of the coastal flow, but a less favorable comparison is obtained with the measurements (the Fourier norm is 1.22).

**Figure A1.**
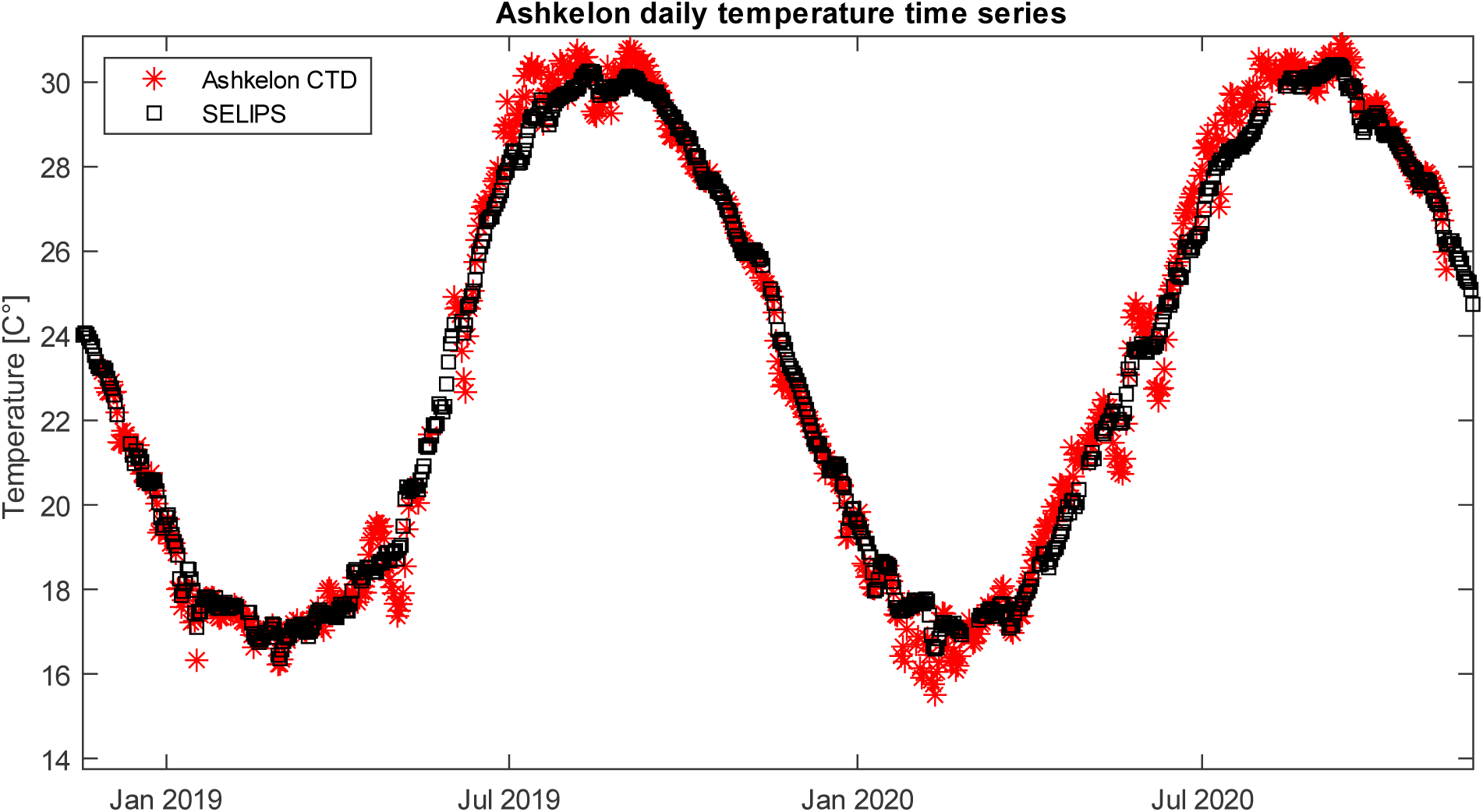
Circadian averages of temperature as measured at the Ashkelon measuring station at a water depth of 11 m (red) against the results of the SELIPS model in the same location. The calculated correlation coefficient between the two time series is 0.99 with an RMSE of 0.55 degrees.

**Figure A2.**
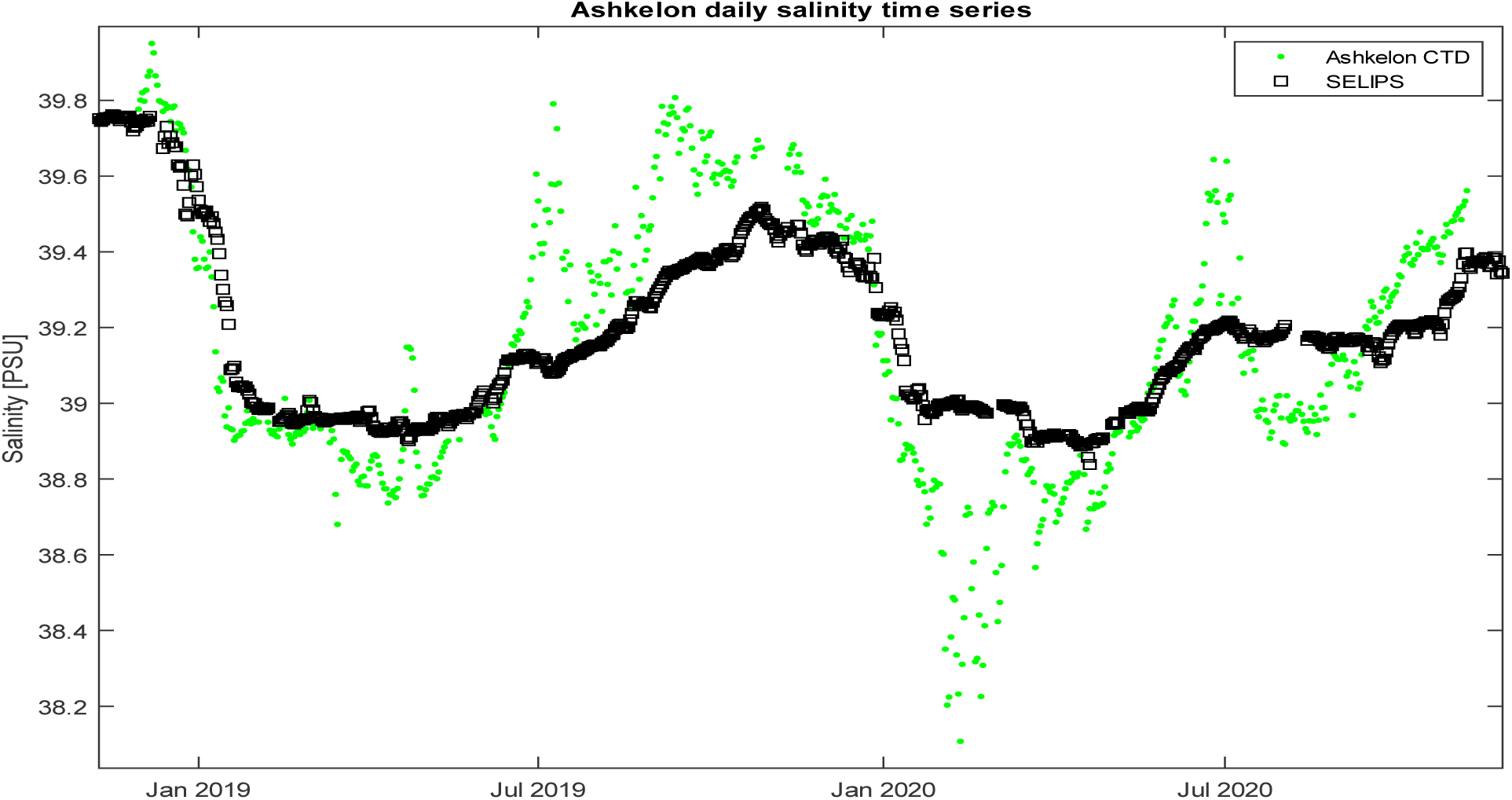
Circadian averages of salinity as measured at the Ashkelon measuring station at a water depth of 11 m (green) against the results of the SELIPS model in the same location. The calculated correlation coefficient between the two-time series is 0.81 with RMSE of 0.2 PSU.

**Figure A3.**
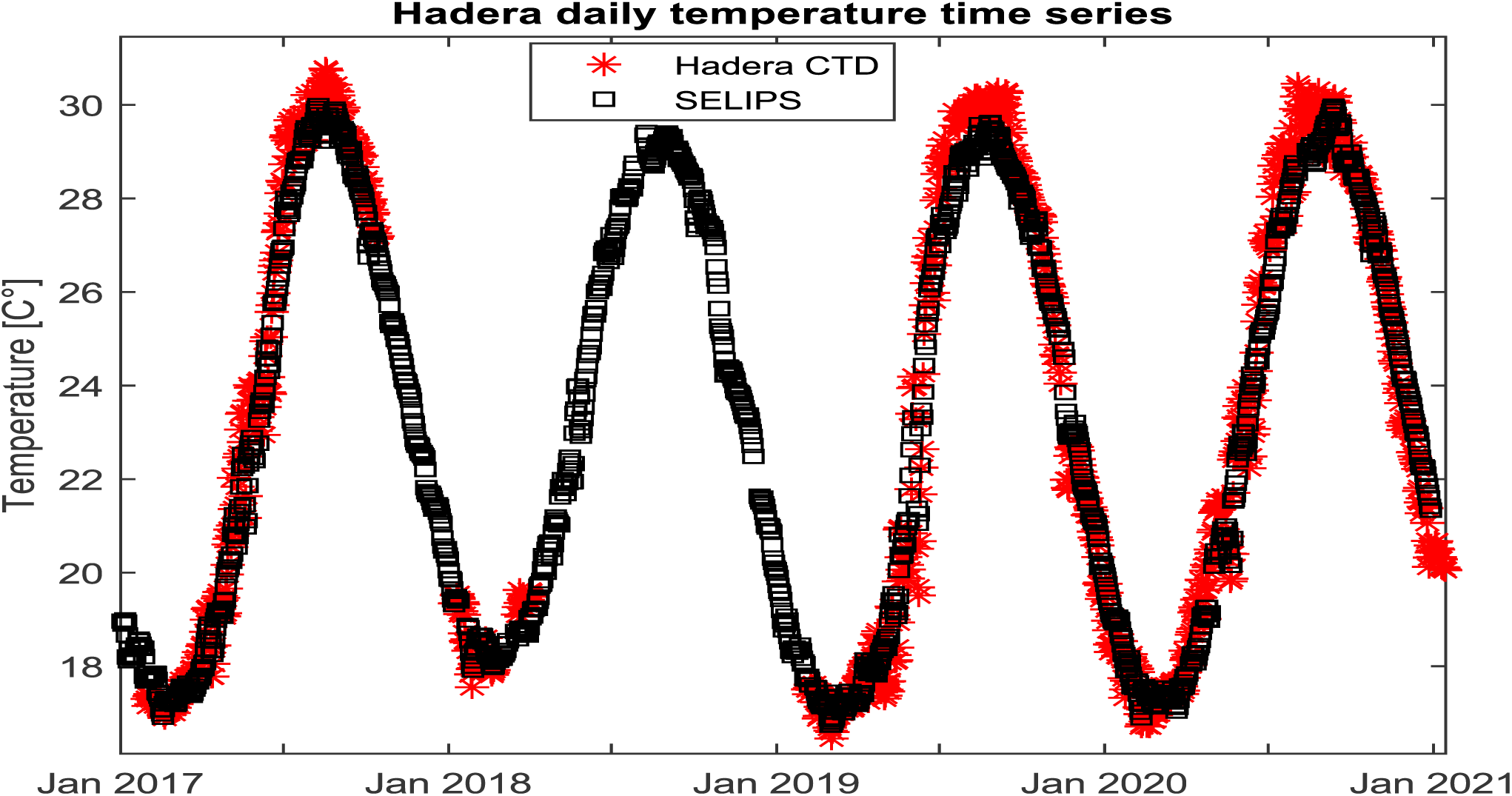
Circadian averages of temperature as measured at the Hadera measuring station at a water depth of 11 m (red) against the results of the SELIPS model in the same location. The calculated correlation coefficient between the two time series is 0.99 with an RMSE of 0.59 degrees.

**Figure A4.**
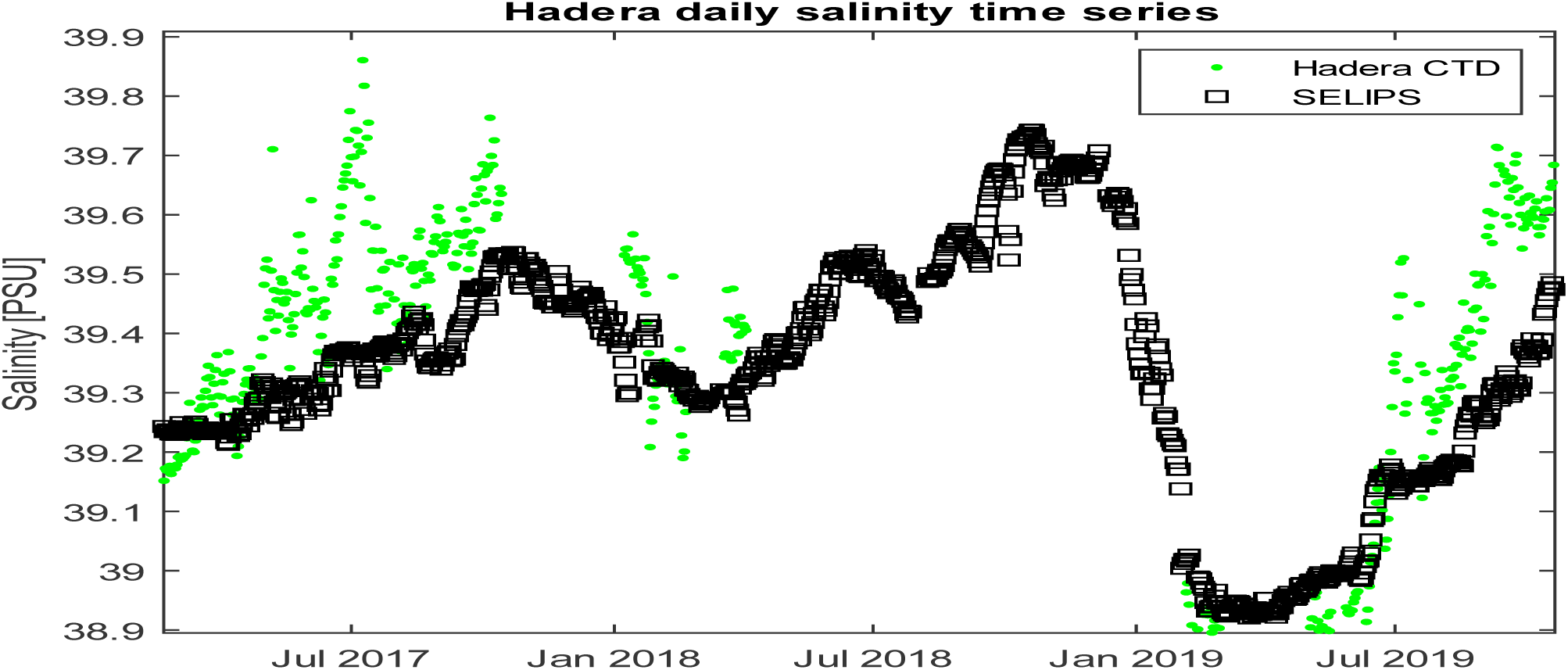
Circadian averages of salinity as measured at the Hadera measuring station at a water depth of 11 m (red) against the results of the SELIPS model in the same location. The calculated correlation coefficient between the two time series is 0. 91 with RMSE of 0.11 PSU.

**Figure A5.**
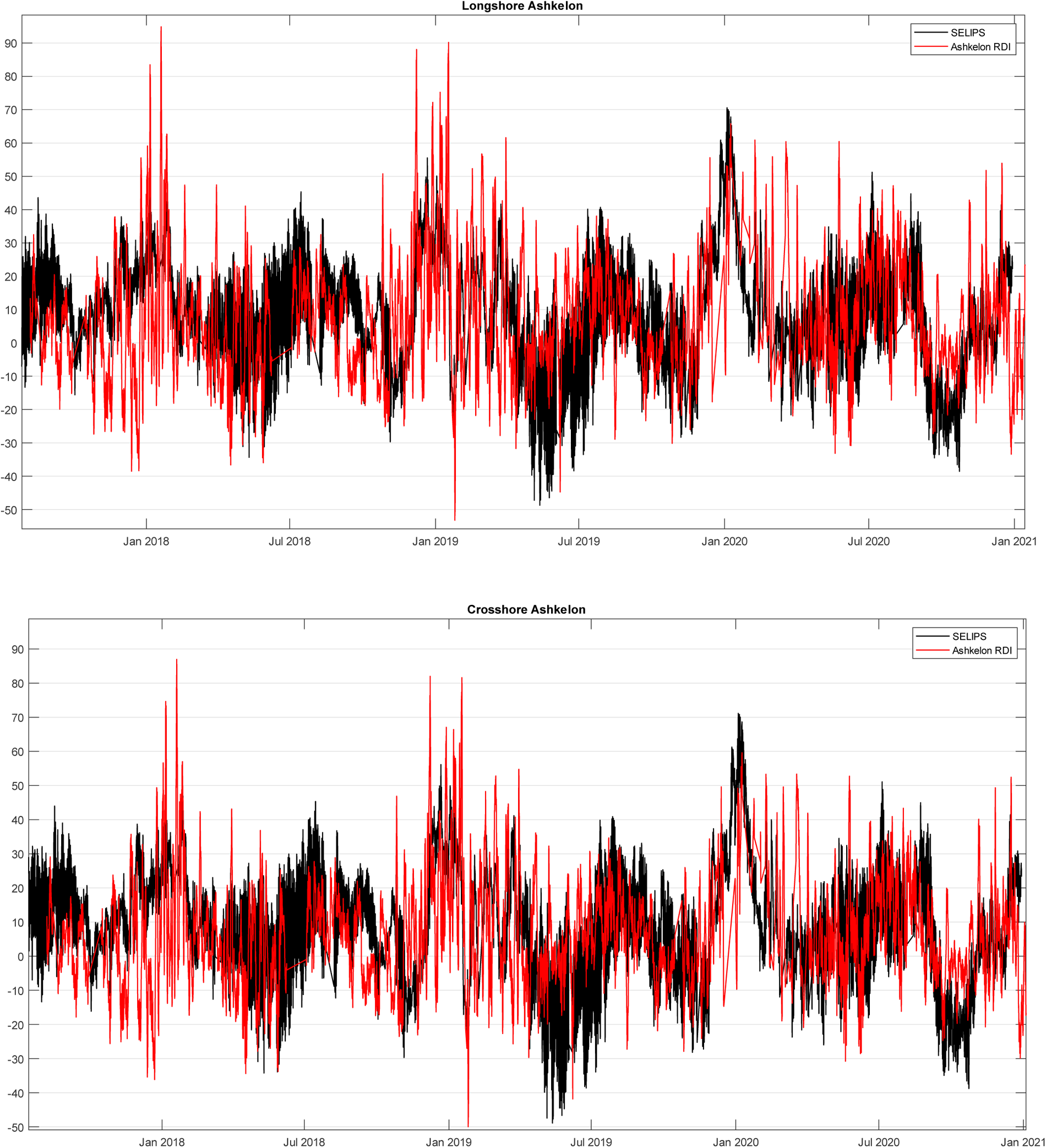
3-hour averages of the components of the currents along (top) and perpendicular (bottom) to the lines with a depth value as measured at the Ashkelon station between 2021 and 207 at a water depth of 11 m (red) against the results of the SELIPS model in the same position (black). The calculated correlation coefficients between the two time series are 0.37 for the tangential component and 0.34 for the component perpendicular to the isobath.

**Figure A6.**
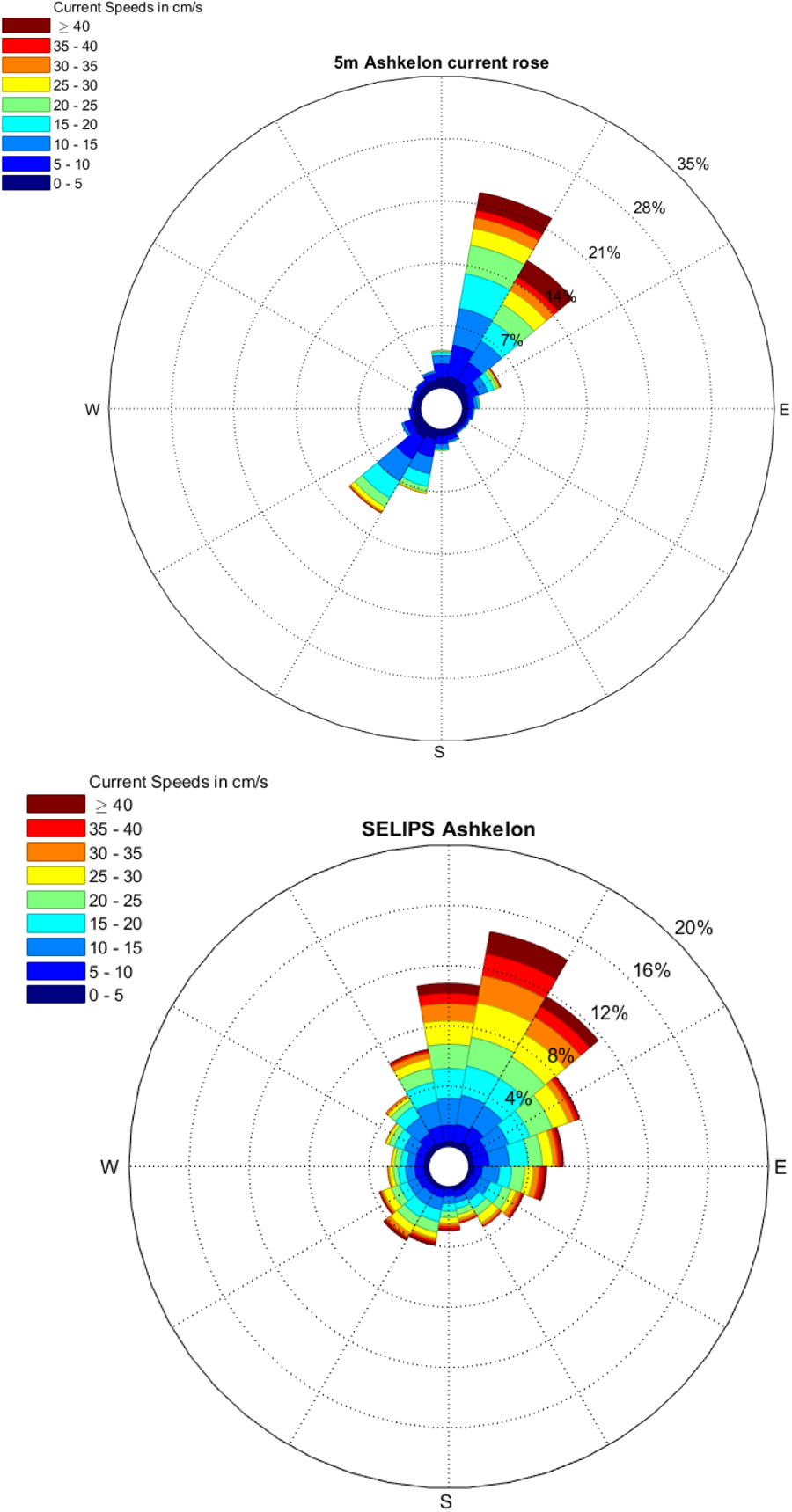
Rose diagram as calculated based on the measurements in Ashkelon at a depth of 5 m (top) for the years 2017-2021 against the results of the SELIPS model in the same location (bottom).

**Figure A7.**
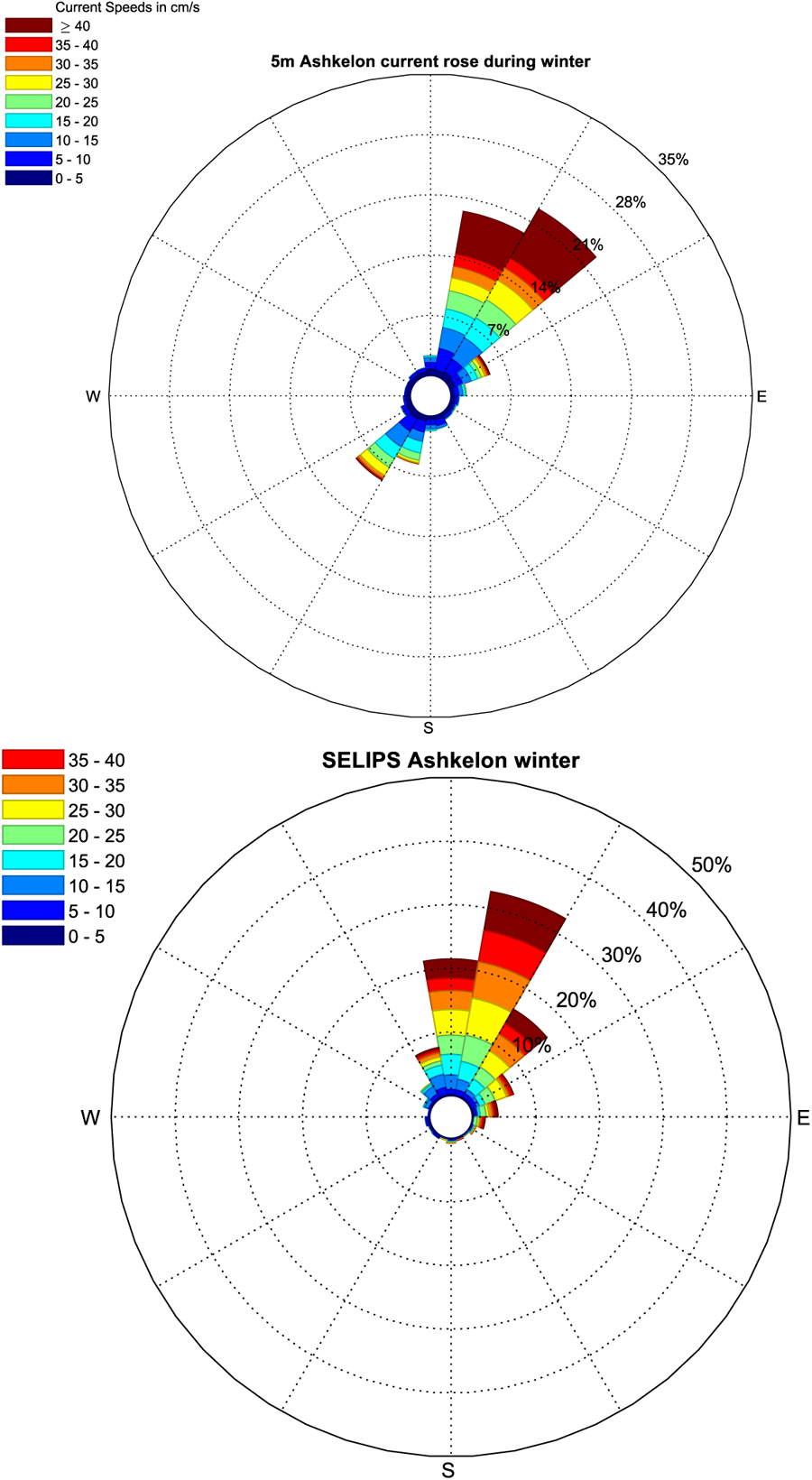
Rose diagram as calculated based on the measurements in Ashkelon at a depth of 5 m (top) in the winters of 2017-2021 against the results of the SELIPS model in the same location (bottom).

**Figure A8.**
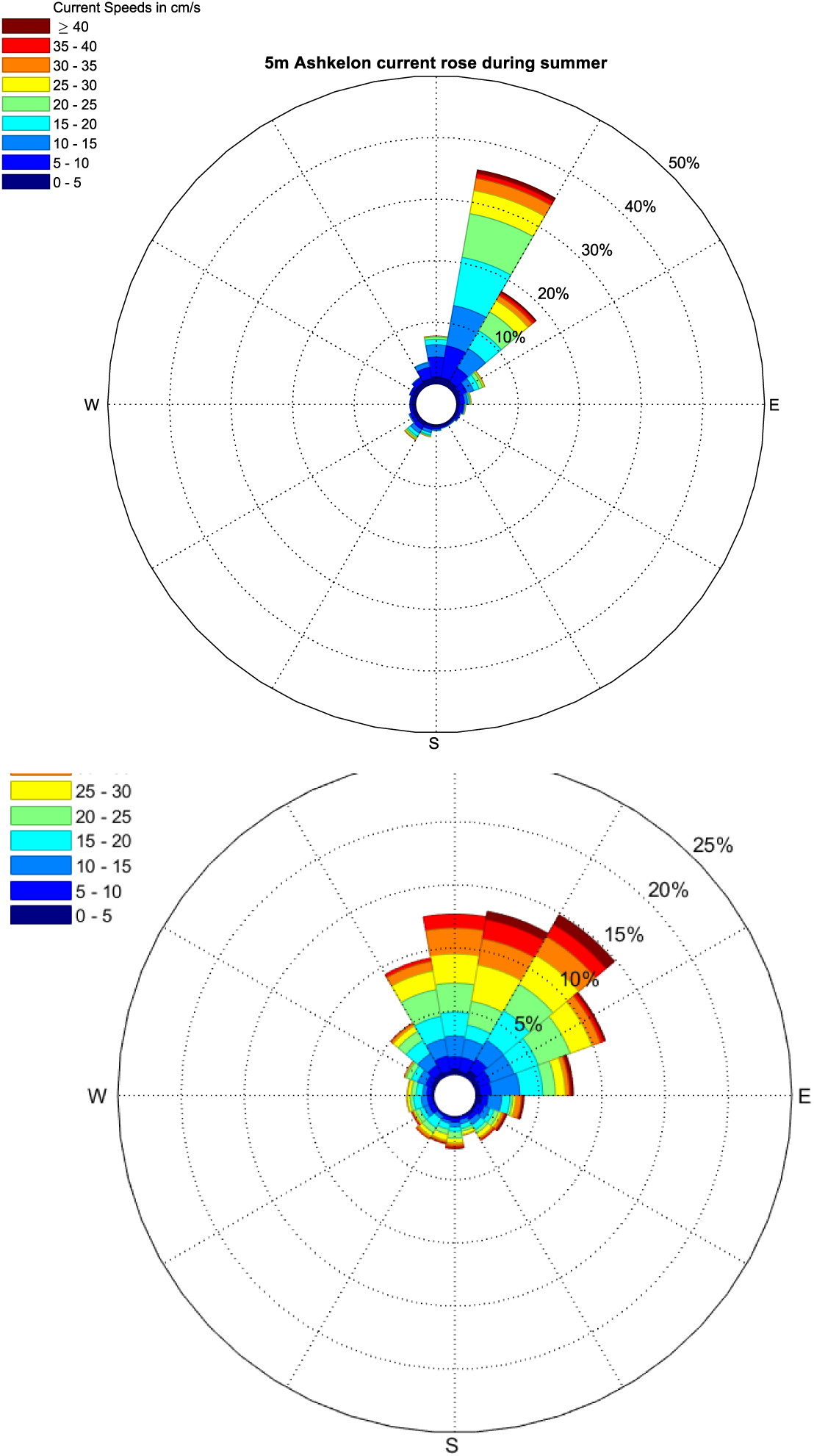
Rose diagram as calculated based on the measurements in Ashkelon at a depth of 5 m (top) in the summers of 2017-2021 against the results of the SELIPS model in the same location (bottom).

**Figure A9.**
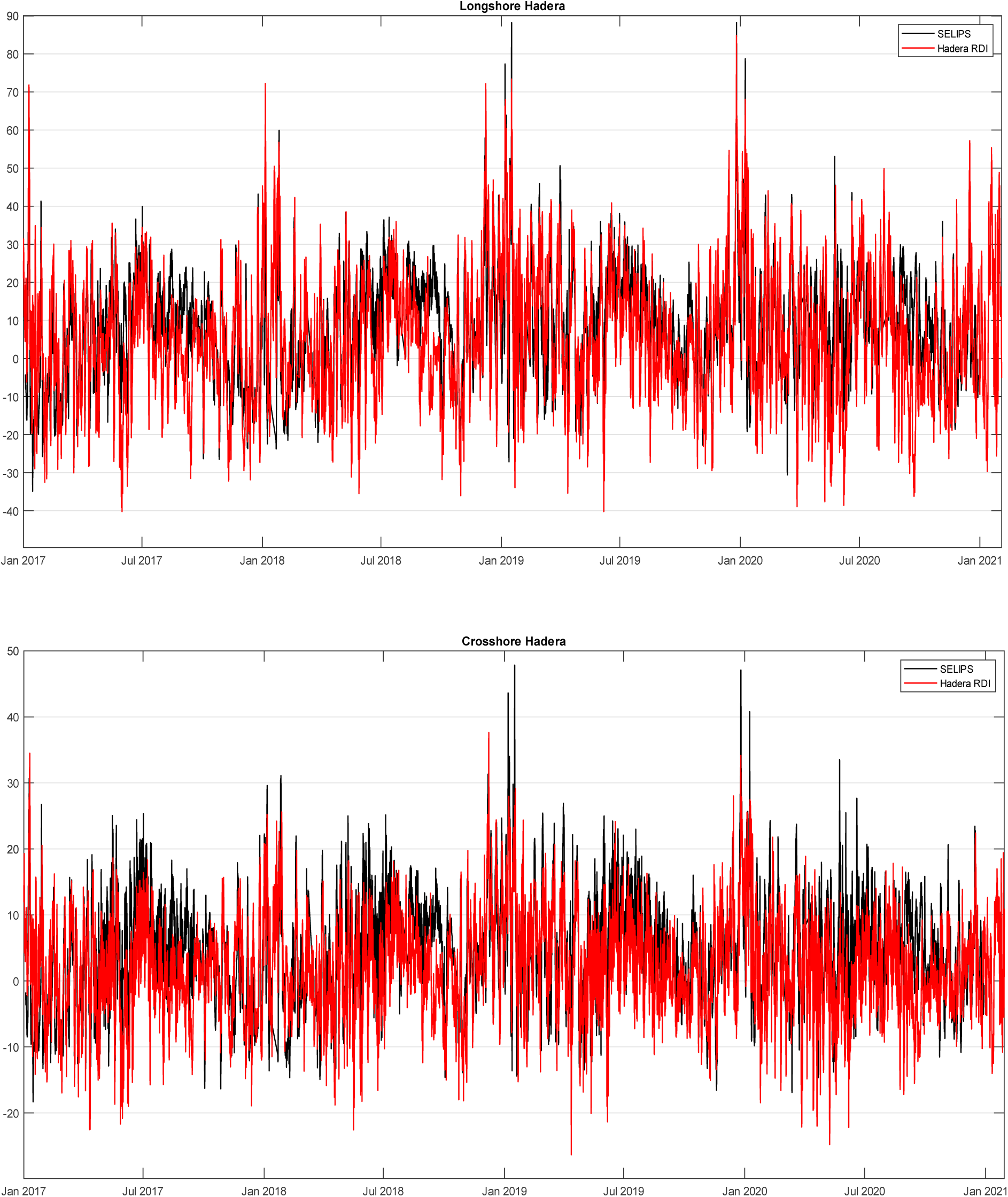
Three-hour averages of the components of the currents along (top) and perpendicular (bottom) to the isobaths as measured at the Hadera station between 2021 and 207 at a water depth of 11 m (red) against the results of the SELIPS model in the same position (black). The calculated correlation coefficients between the two time series are 0.67 for the tangential component and 0.61 for the component perpendicular to the bank depth line.

**Figure A10.**
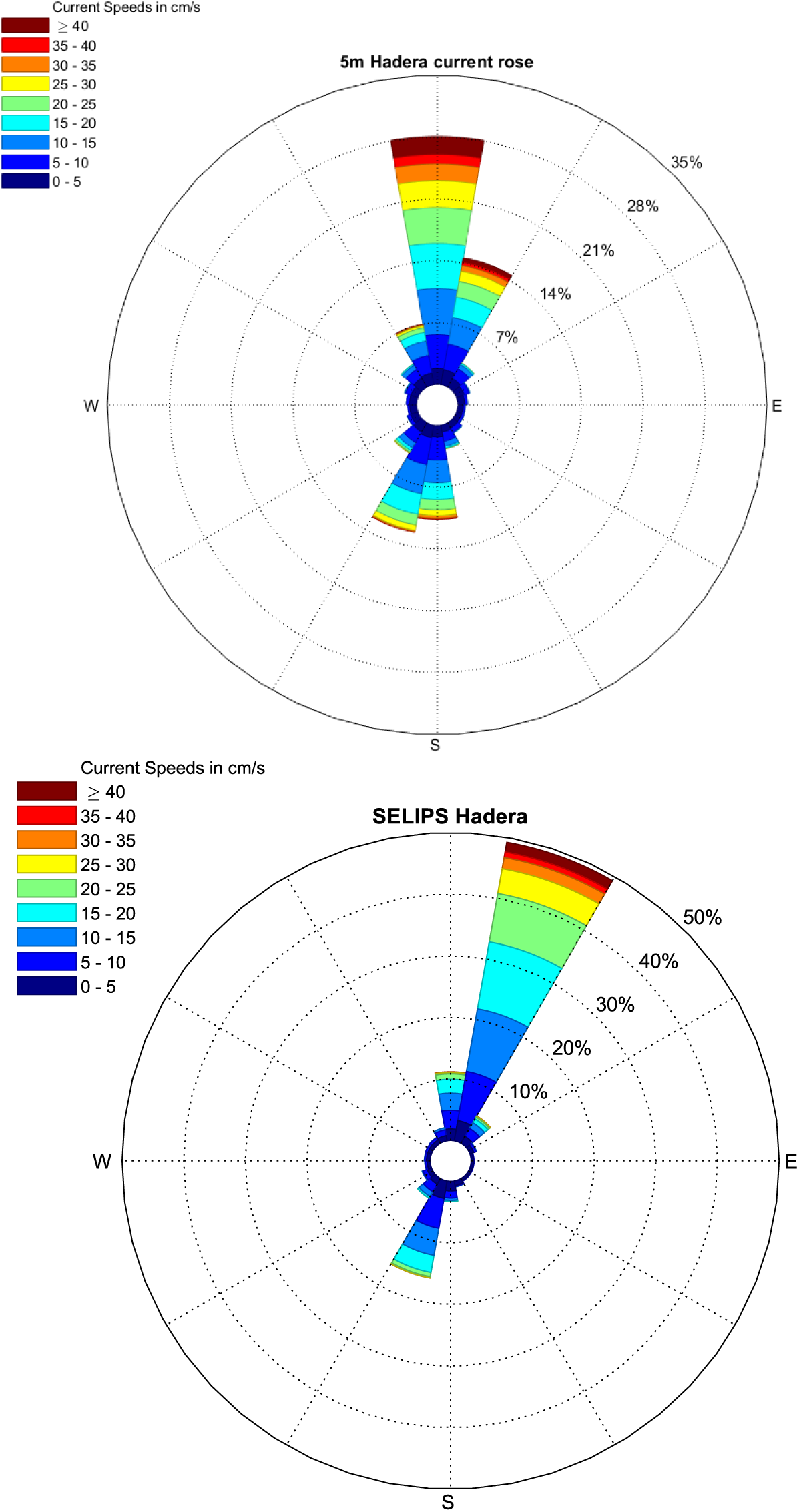
Rose diagrams as calculated based on the measurements in Hadera at a depth of 5 m (top) for the years 2017-2021 against the results of the SELIPS model in the same location (bottom).

**Figure A11.**
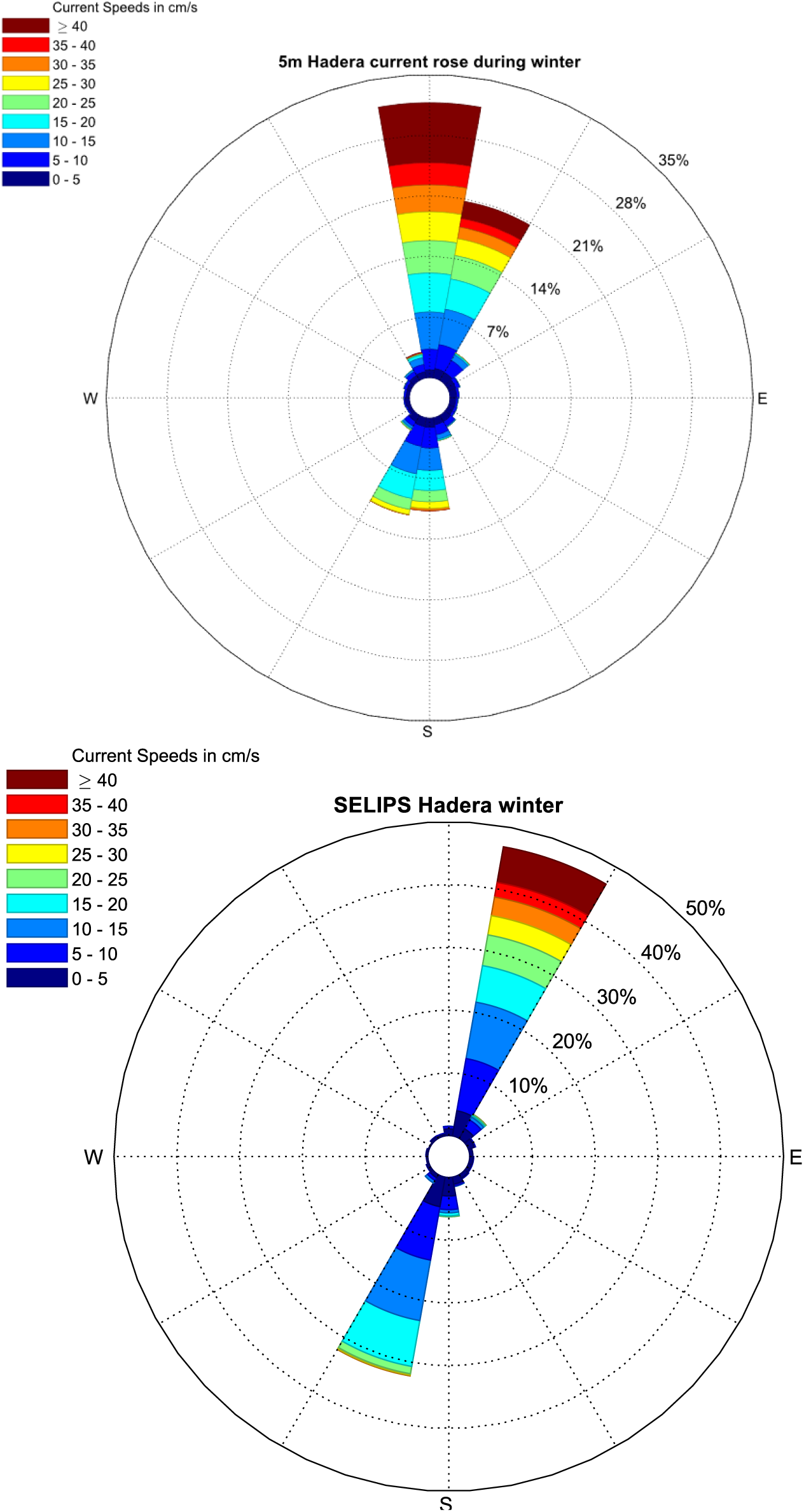
Rose diagrams as calculated based on the measurements in Hadera at a depth of 5 m (top) in the winters of 2017-2021 against the results of the SELIPS model in the same location (bottom).

**Figure A12.**
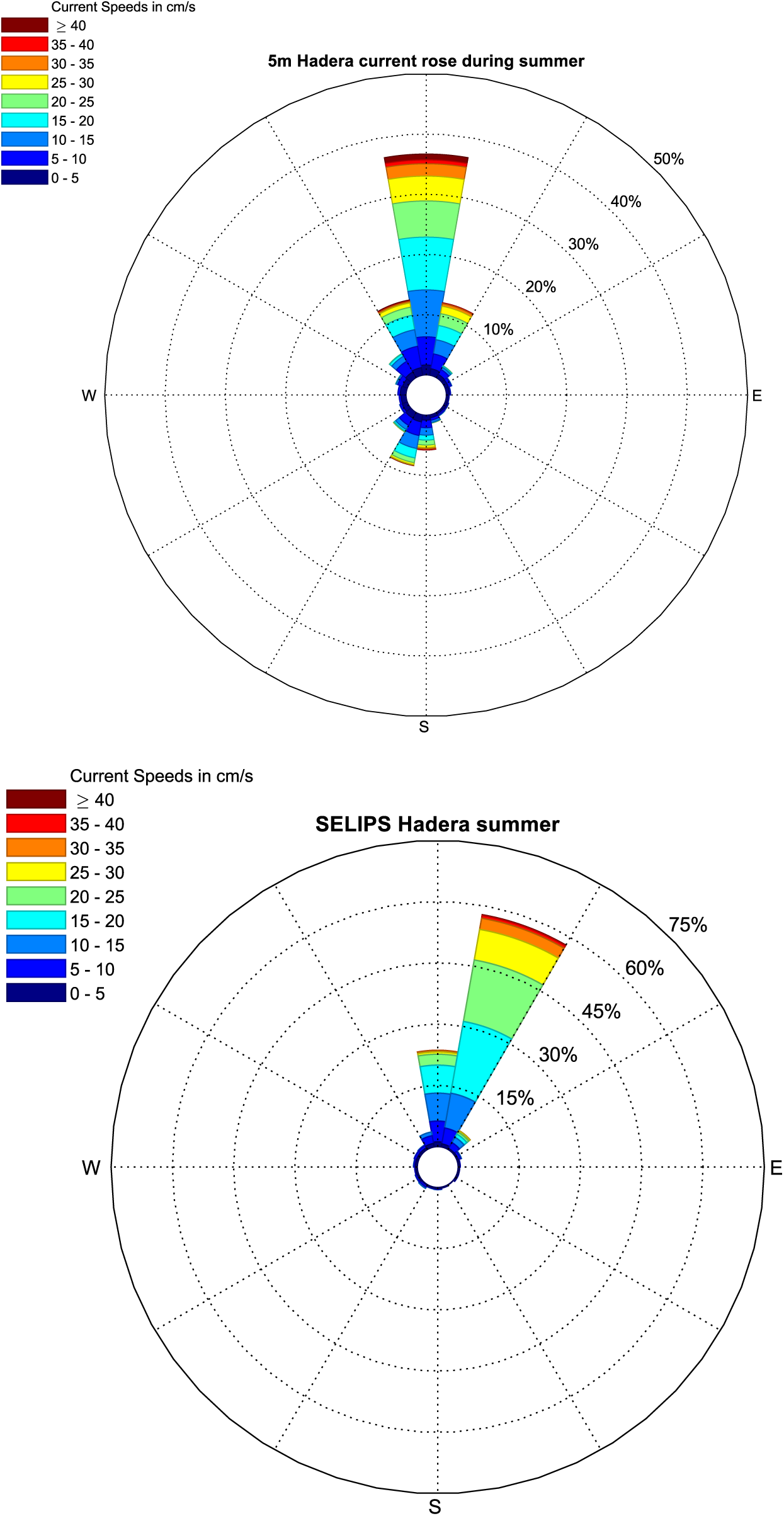
Rose diagrams as calculated based on the measurements in Hadera at a depth of 5 m (top) in the summers of 2017-2021 against the results of the SELIPS model in the same location (bottom).

##### The comparison with the DEEPLEV deep sea sampling station

DEEPLEV is a marine monitoring station located in the open sea (32° 59′ 58.2000“ N, 34° 29′ 58.8120” E) about 50 km off the northern coast of Israel, where data on sea currents and hydrographic conditions are collected along the water column. The results of the model were compared with the temperature data measured at a depth of 30 meters, and with the data of the currents measured at a depth of 50 meters. All observations were proposed every three hours in order to coincide with the times of the model results. In general, a good comparison is obtained between the results of the model and the temperature values measured at a depth of 30, although there is an underestimation of the hot temperatures in the summer season (Fig. A13). A comparison of the rose diagrams of the currents at a depth of 50 meters shows that statistically the model manages to model the general movement towards the north, but the movement towards it east and south is less dominant than those observed. The flow velocity in the model is slightly lower than the measured flow in all directions (Fig. A14).

**Figure A13:**
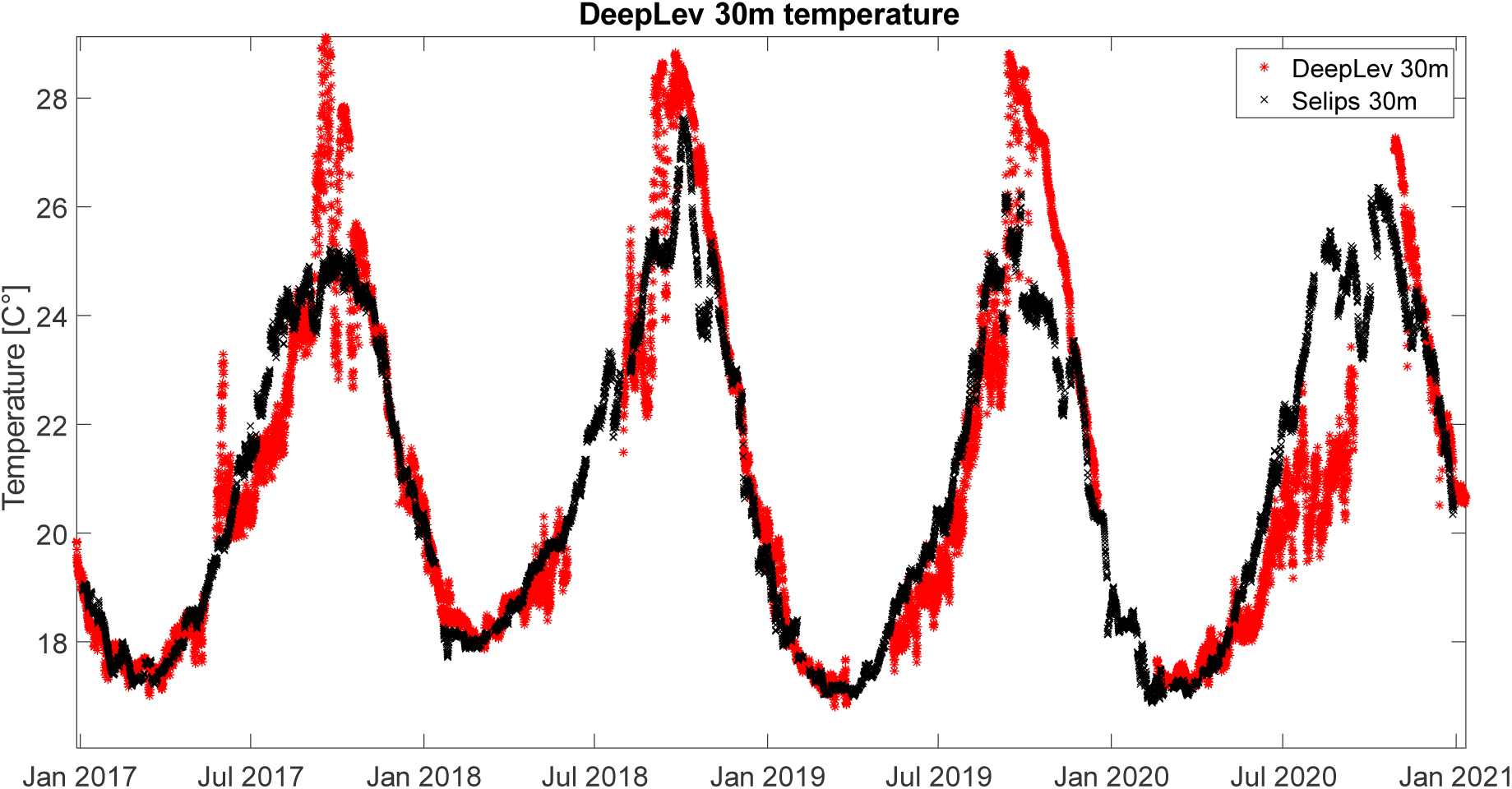
Temperature measured at the DEEPLEV measuring station at a water depth of 30 m (red) against the results of the SELIPS model in the same position (black). The calculated correlation coefficient between the two time series is 0.9 1 with an RMSE of 1.38°C.

**Figure A14.**
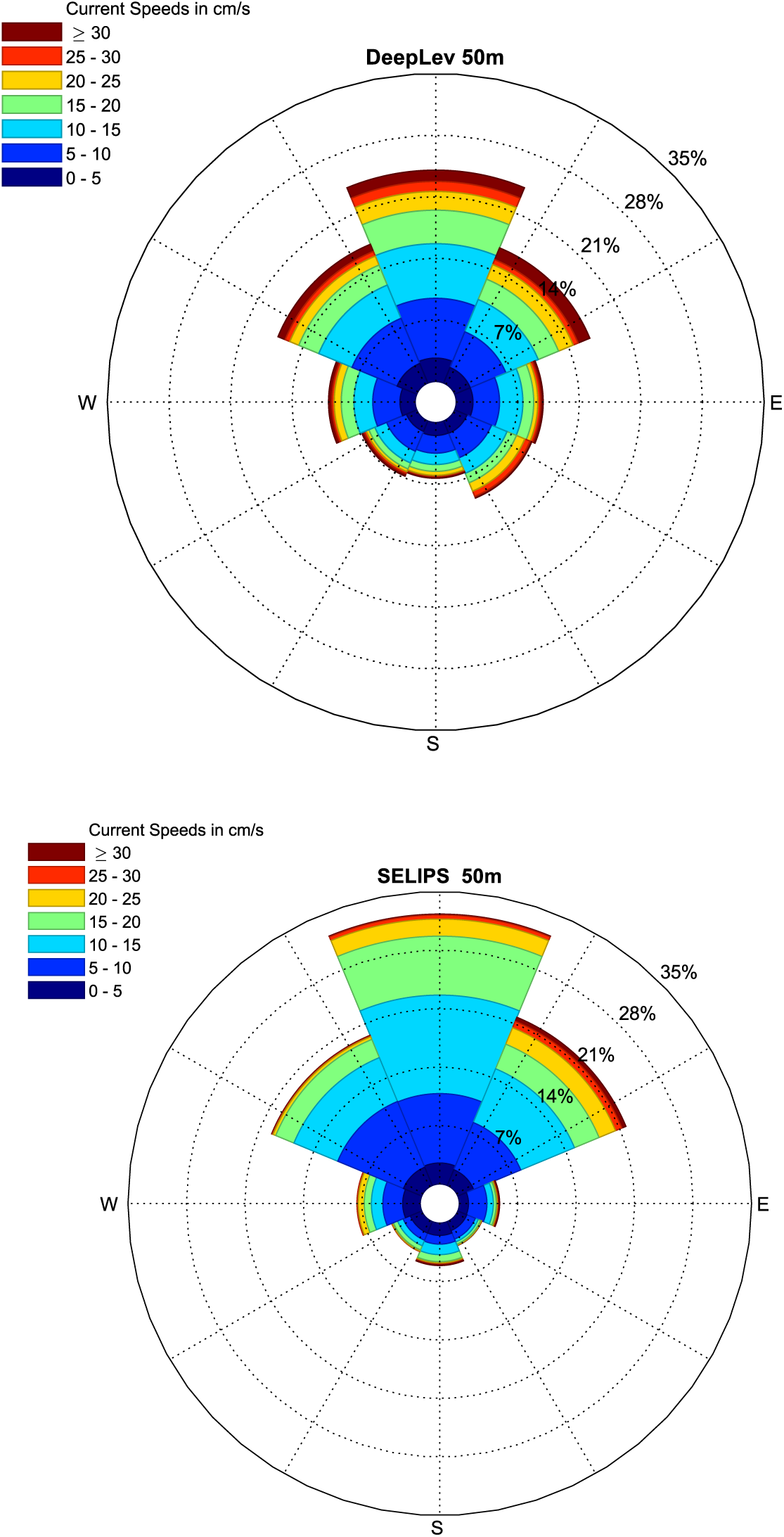
Rose diagram as calculated based on the measurements at the DEEPLEV deep sea sampling station at a depth of 50 m (top) against the results of the SELIPS model in the same location (bottom).

### Supplementary section S2: Biological component elaborated results and discussion

#### Elaborated biological methods

Each site was sampled at three stations parallel to the coastline, with depths of 10, 25, and 50 meters. To maximize catch per cruise, three sampling methods were used. (A) Plankton light trap (Bellamare, USA), which were deployed at dusk and collected before dawn. Light traps were steadily placed two meters below the water surface; (B) Bongo net (Ocean Instruments Ltd, USA) - with a mesh size of 300µm, filtering the mid-layer of the water column; (C) Manta net (Ocean Instruments Ltd, USA), sampling the top layer of the water (Neuston) with a mesh size of 300µm and an opening of 1.275 m². Bongo and Manta nets were towed for 20 min at speed of 2 knots by a small boat and were mounted with a mechanical Flow meter (Hydro-Bios, Germany) to standardize the filtered water volumes. All samples were sieved on board through a 200µm sieve and immediately preserved in Absolute ethanol for further analysis in the lab. Each sampling cruise contained nine samples of Manta, Bongo and light trap (three for each).

In the lab, fish eggs and larvae were separated from the total catch into different microtubes and counted using a light stereo microscope Stemi 508 (Zeiss, Germany). Total genomic DNA was extracted using the AccuPrep® Genomic DNA Extraction Kit (BIONEER, USA) according to the manufacturer’s specifications

The following protocol was used to identify the larvae. First, morphologically unique larvae were picked arbitrary from samples (FAO, leis book), imaged using a stereoscope-mounted camera Axiocam 105 Color (Zeiss, Germany) and subjected to molecular identification. Next, extracted DNA was subjected to DNA barcoding methodology, as follows. PCR amplifications were performed for the commonly used cytochrome c oxidase subunit I gene (COI) using the primers and protocols of (Ward et al., 2005). Obtained PCR products were verified on gel and sent to sequencing at HyLaboratories Ltd. (Rehovot, Israel). Multispecies metabarcoding approach

Extracted total genomic DNA of pooled egg and larvae samples was subjected to metabarcoding methodology. The mitochondrial 12S rRNA gene was amplified following the primers amended with SP1/SP2 tags and protocols of (Miya et al., 2015). Obtained PCR products were sent to Syntezza Bioscience Ltd. (Jerusalem, Israel) for library preparation and sequencing of 2x250 bp Illumina MiSeq reads.

Demultiplexed paired-end reads were processed with R v. 3.6.3 (R Core Team, 2021) package “biostring” and “DADA2” pipeline (Callahan et al., 2016; Pages et al., 2013). Primer sequences were trimmed using “biostring” and then, DADA2 was used for sequence assembly, quality filtration, and chimera removal. Obtained amplicon sequence variants (ASVs) were compared against NCBI and BOLD databases for specific identification. Ambiguous results were classified to family level or declared as “unknown”.

Final sequences of both barcoding and metabarcoding were compared online via NCBI (https://www.ncbi.nlm.nih.gov/) and BOLD (https://boldsystems.org/) databases, considering a ≥ 98% genetic similarity as a valid species-level confirmation.

Last, univariate statistical analyses of non-parametric tests (Mann-Whitney, Kruskal-Wallis) have been performed using SPSS ver. 20 (IBM). Multivariate analyses such as test for Analysis of Similarity (ANOSIM) and Similarity Percentage (SIMPER) have been performed using PRIMER ver.5.2.9 (PRIMER-E).

### Elaborated biological results

Throughout the sampling period a total of 48 Manta trawls, 48 Bongo trawls and 52 light traps were collected throughout the sampling period of August 2019 to August 2021. Out of which only the samples from August 2019 to November 2020 (n=39) were processed due to underestimated workload. Total counting of all samples yielded 84804 eggs and 7370 larvae in total.

**Table B1:** Cruise dates per site.

| Achziv | Shikmona | Gdor | Avtah |
| --- | --- | --- | --- |
| 16/08/2019 | 10/06/2020 | 26/08/2020 | 11/05/2020 |
| 20/05/2020 | 07/07/2020 | 02/11/2020* | 28/07/2020 |
| 30/06/2021 | 02/05/2021 | 29/06/2021 | 23/06/2021 |
| 01/08/2021 | 07/07/2021 | 24/08/2021 | 11/08/2021 |

\* Gdor cruise of 02/11/2020 was conducted in November due to COVID-19 restrictions.

### Mean density of ichthyoplankton

Mean density (individual/m^3^) of fish eggs and larvae were compared for three parameters: (1) water depth of 10, 25, and 50 meters; (2) position of sample in water column (surface ver. middle); (3) and sampling sites. No significant difference in mean density of fish eggs was found between water depth (Kruskal-Wallis, p=0.886) and water layer (Mann-Whitney, p=0.606). Comparison between the sites did show a significant difference (Kruskal–Wallis, p<0.05) with Achziv and Shikmona sites showing the highest fish eggs mean density of 16 ind/m^3^ and 7.36 ind/m^3^ respectively (Fig B1). No significant differences in fish larvae density were found between water depth (Kruskal-Wallis, p=0.838), water layer (Mann-Whitney, p=0.349), and sites (Kruskal-Wallis, p=0.103). However, slightly higher fish larvae densities were found in the northern sites of Achziv and Shikmona with mean density of 0.942 ind/m^3^ and 0.931 ind/m^3^ respectively (Fig B2)

**Figure B1:**
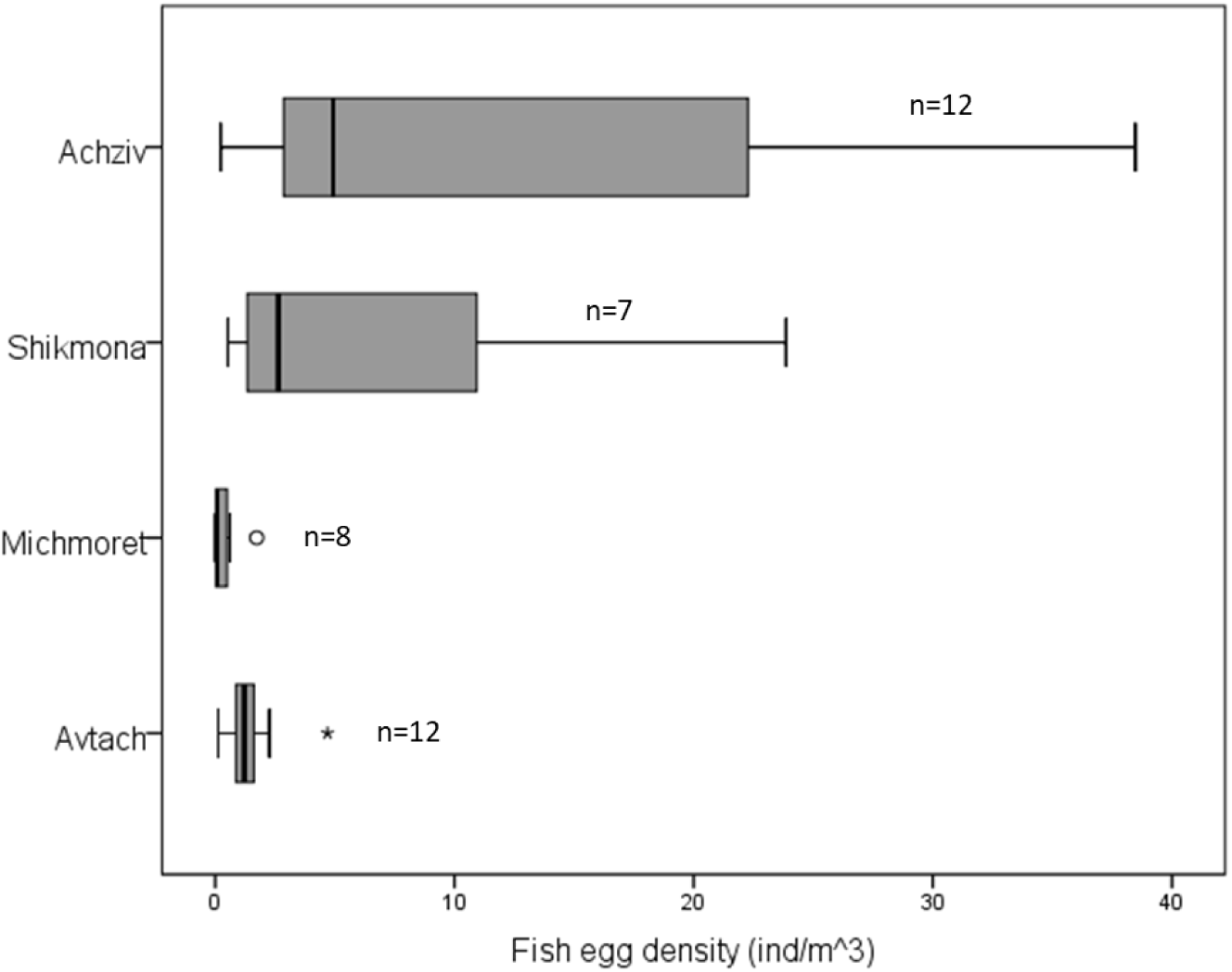
Comparison of fish egg density between the four sites, organized from north to south.

**Figure B2:**
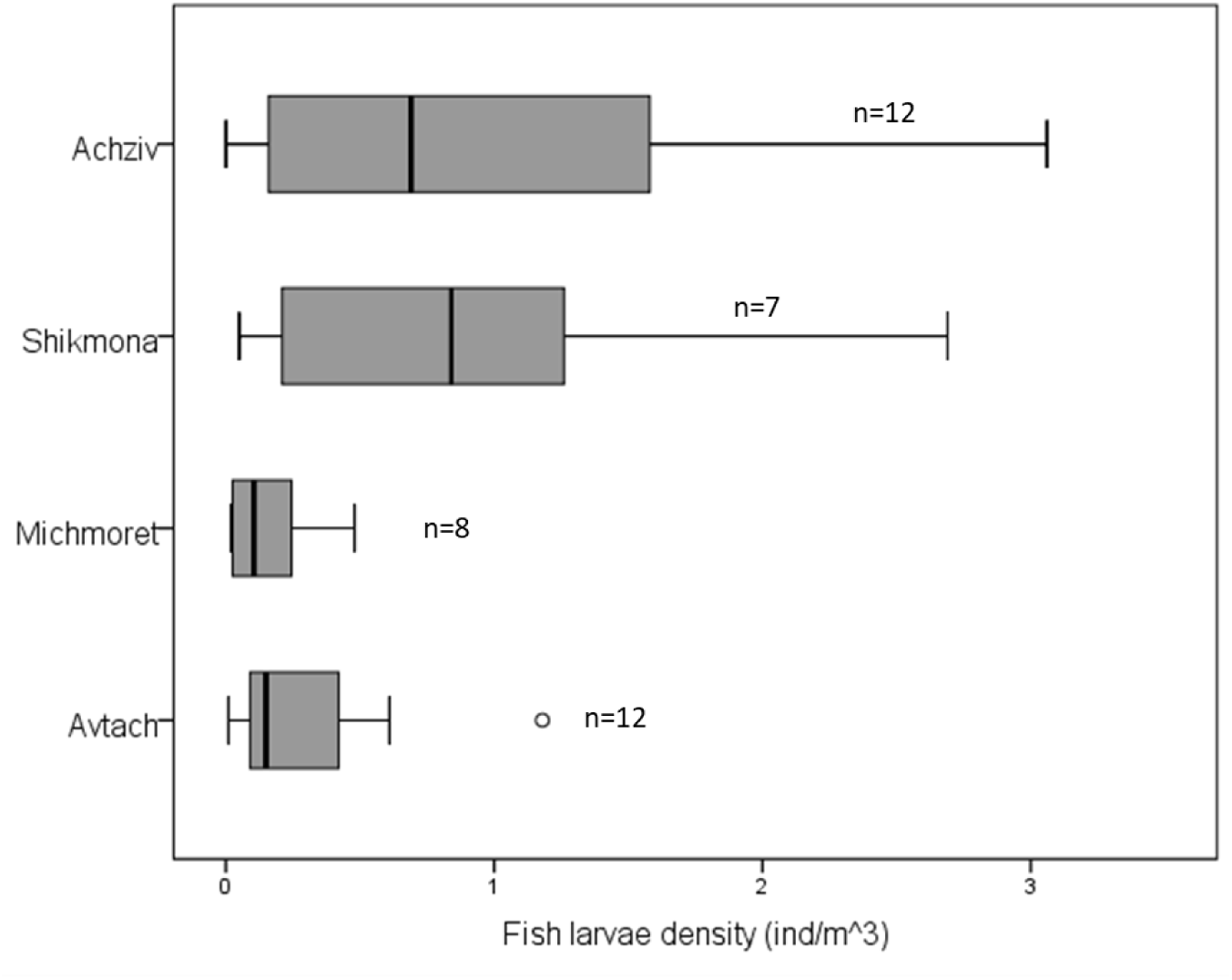
Comparison of fish larvae density between the four sites, organized from north to south.

### Species composition

In an ANOSIM analysis, no significant difference was found in species composition of fish eggs between the sites (R=0.118), water depth stations (R=0.044) and water layer (R=-0.051). Non-significant differences were also found in larval species composition (R=0.111, R=0.042, R=0.011), respectively. In accordance with the habitat structure, rocky grouper from the genus *Epinephelus* were more abundant in Achziv, and pelagic species such as *Pomatomus saltatrix, Scomberomorus commerson* and *Decapterus russelli* were more abundant in the southern sandy habitat of Avtach (Table B3).

Pooled data shows that the most prevalent species found

**Figure B3:**
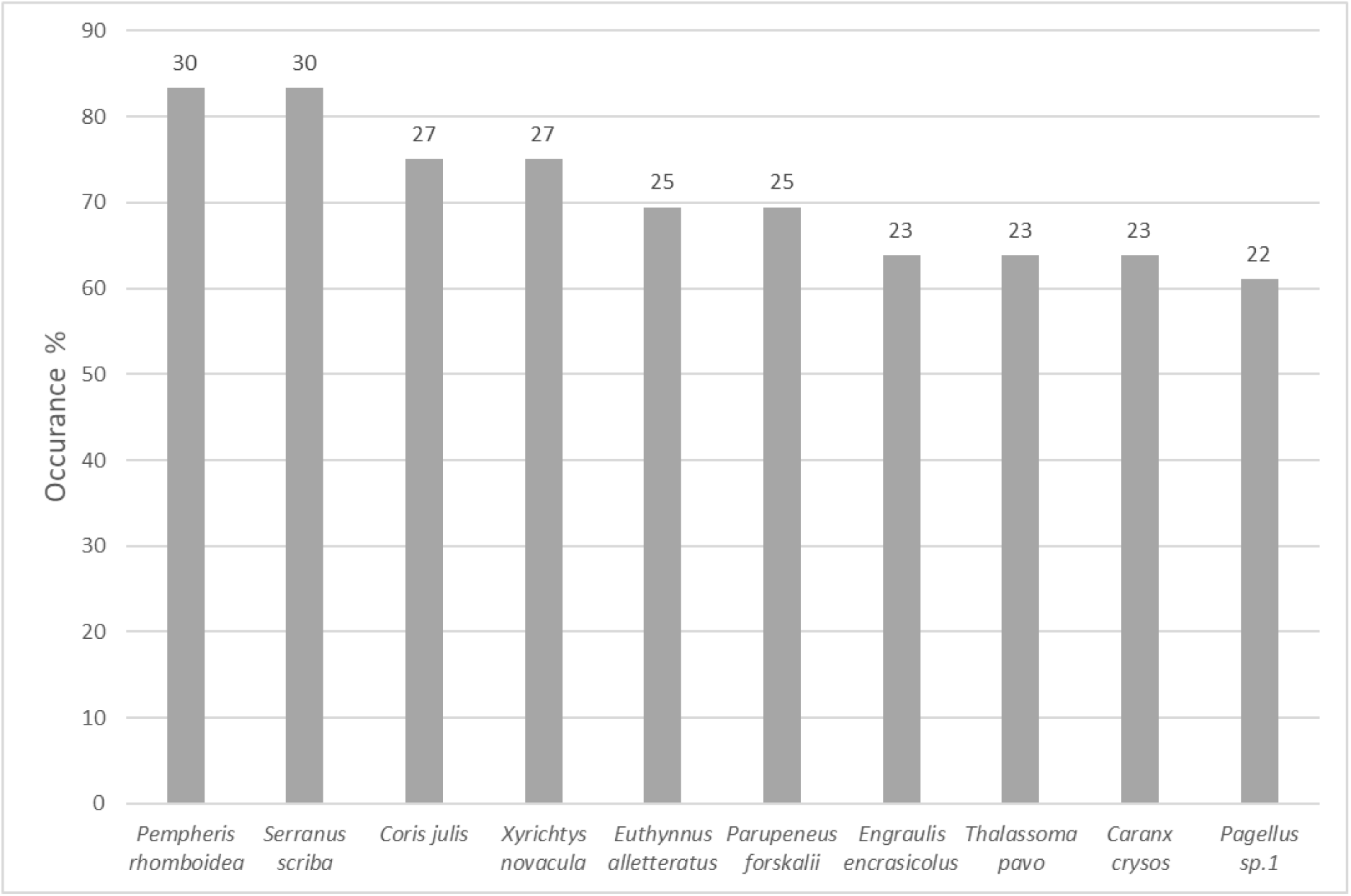
Fish <u>eggs</u> occurrences of the ten most dominant species. Number on top of columns indicate the number of times a species was found in samples, based on presence only data. Samples n = 36.

**Figure B4:**
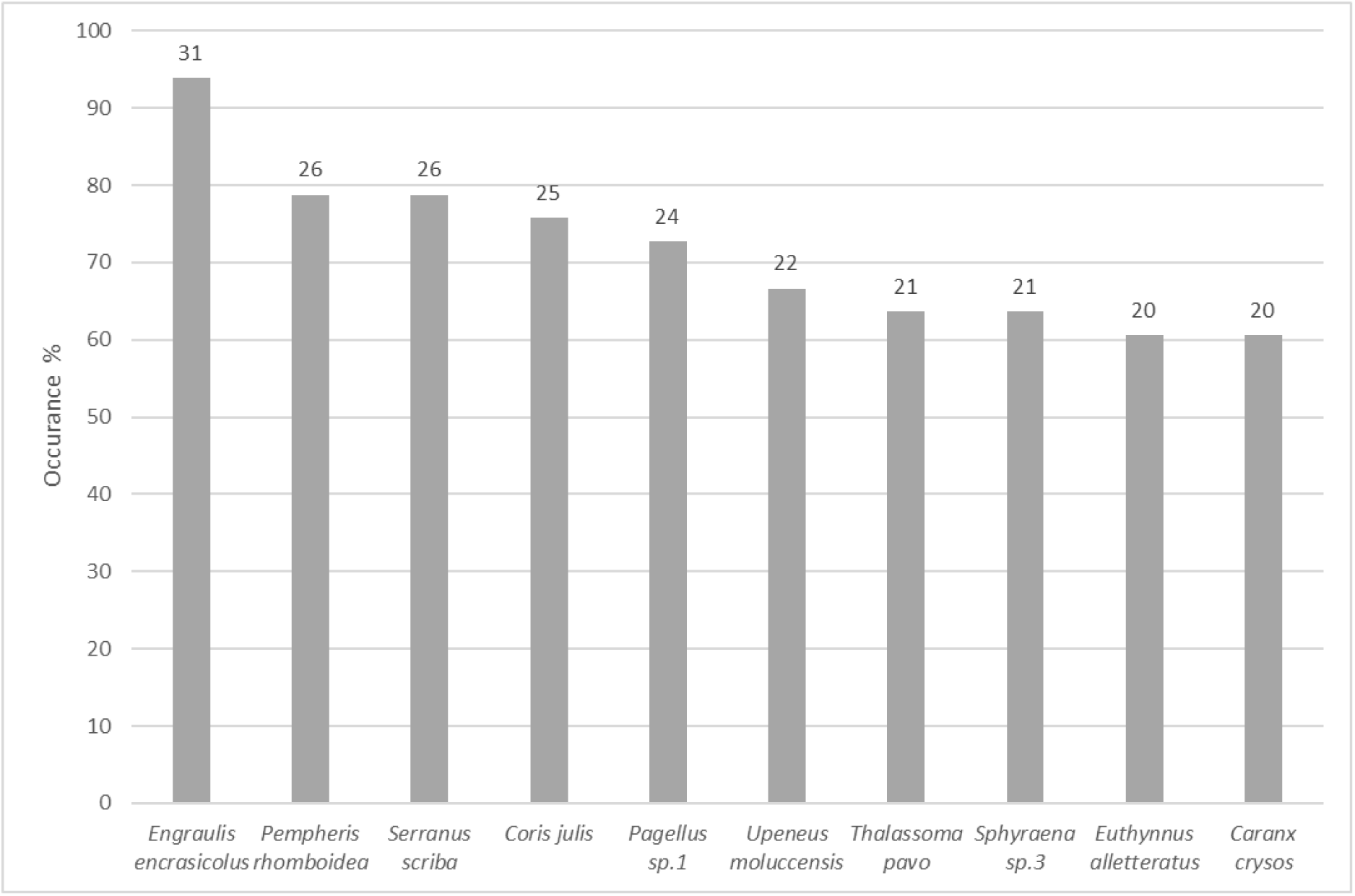
Fish <u>larvae</u> occurrences of the ten most dominant species. Numbers on top of columns indicate the number of times a species was found in samples based on presence only data. Samples n = 33.

### Non-metric multidimensional scaling (nMDS) analysis

nMDS analyses indicate that there is no clustering among sites, north-south gradient, sampling methods, and depth gradient, suggesting that the larval pool is well mixed, or at least the dominant larvae are common across these gradients.

**Figure B5:**
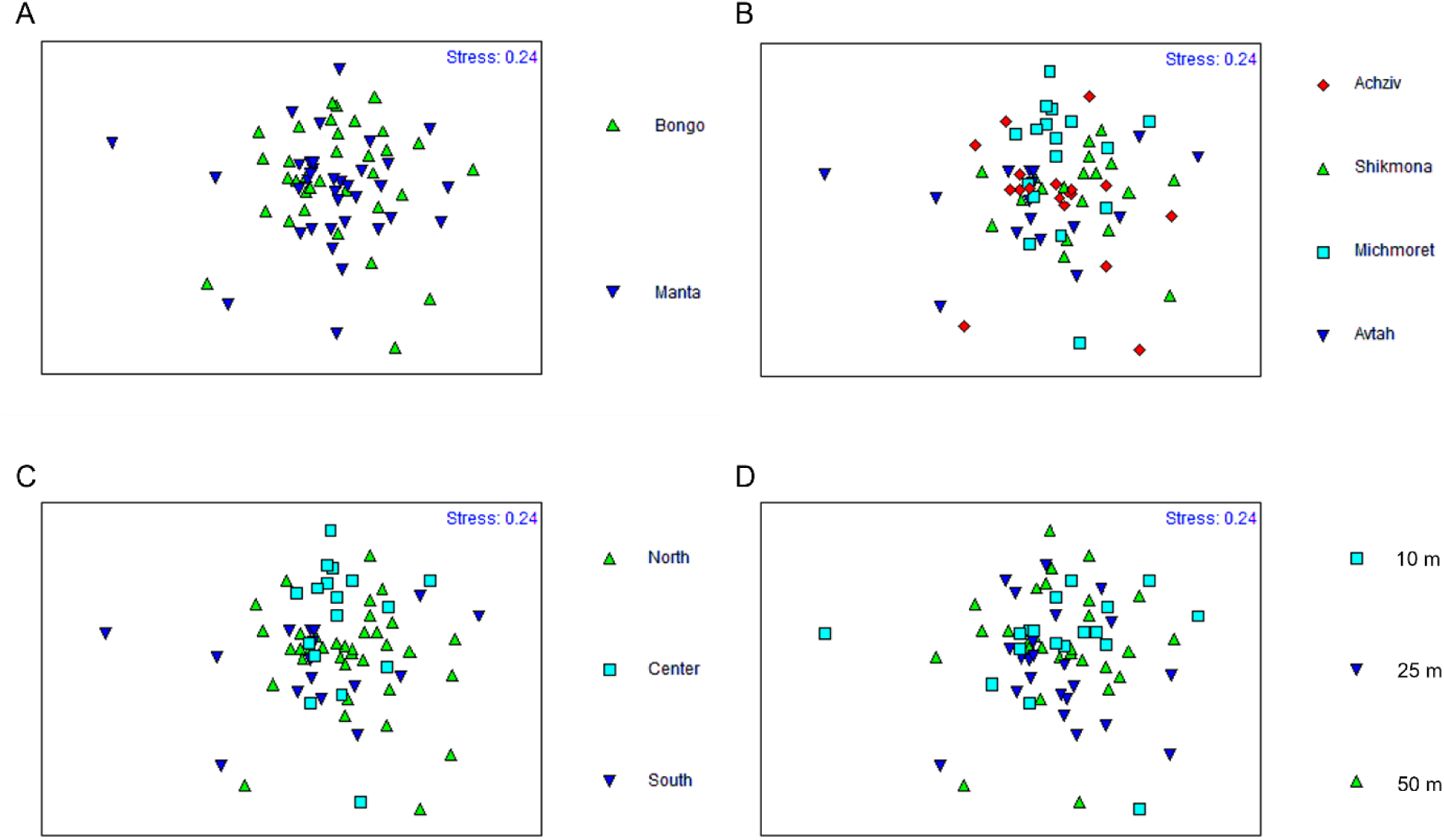
Non-metric multidimensional scaling (nMDS) analysis of the ichthyoplankton samples based on: (A) sampling method, (B) sampling site, (C) north-south gradient, and (D) bottom depth.

### Species identification

Using molecular identification based on 12S mitochondrial rRNA and COI mitochondrial DNA we have identified 57 different families with 144 different species. Among these, 12 species were identified up to genus level, 14 up to family level, and one unknown specimen without any resemblance to known family level in online databases. The ten most prevalent fish eggs and larvae are given in Fig B1 and B2, respectively, and the complete list of the identified ichthyoplankton is given in Table B2.

#### Invasive species

Out of 117 fully identified species 47 are considered invasive to the Mediterranean. An unrecorded species of gobi *Acentrogobius pflaumii* has been sampled during this study and its report is now under review. In addition, second observations of the non-indigenous species *Ambassis dussumieri* and *Chaetodon austriacus* have been reported during this study, evidencing their population status.

#### Rare and cryptic species

The cryptic gobi *Milleriogobius macrocephalus* that has been last observed in the Israeli Mediterranean in the 1980’s has been colleced during this study, as well as the blackspot seabream *Pagellus bograveo* that exists in the Israeli checklist of native fishes but been caught only once every several years.

#### Bathyal species

Seven fish species that inhabit the edge of the continental shelf, the continental slope and the bathyal plataue were observed during the study. The species are *Ceratoscopelus maderensis, Chlorophthalmus agassizi, Helicolenus dactylopterus, Lepidorhombus whiffiagonis, Lesueurigobius suerii, Trypauchen vagina* and *Stomias boa*.

#### Compatibility with known spawning seasons

Revisiting the spawning seasonality through the collected fish eggs of this study provided new information on three species that their eggs have been collected outside their known spawning season (Table B2), and five species that had no previous knowledge on their spawning season (Table B3).

**Table B2.** Fish eggs findings of species outside their known spawning season.

| Species | Known spawning season | No. of samples in this study |  |  |  |  |
| --- | --- | --- | --- | --- | --- | --- |
|  |  | May | June | July | August | November |
| *Diplodus puntazzo* | September - December | 4 | 1 | 1 | 2 | 3 |
| *Diplodus sargus* | January - March | 5 |  | 2 | 1 |  |
| *Epinephelus costae* | December - May | 8 | 3 | 1 |  | 1 |

**Table B3.**
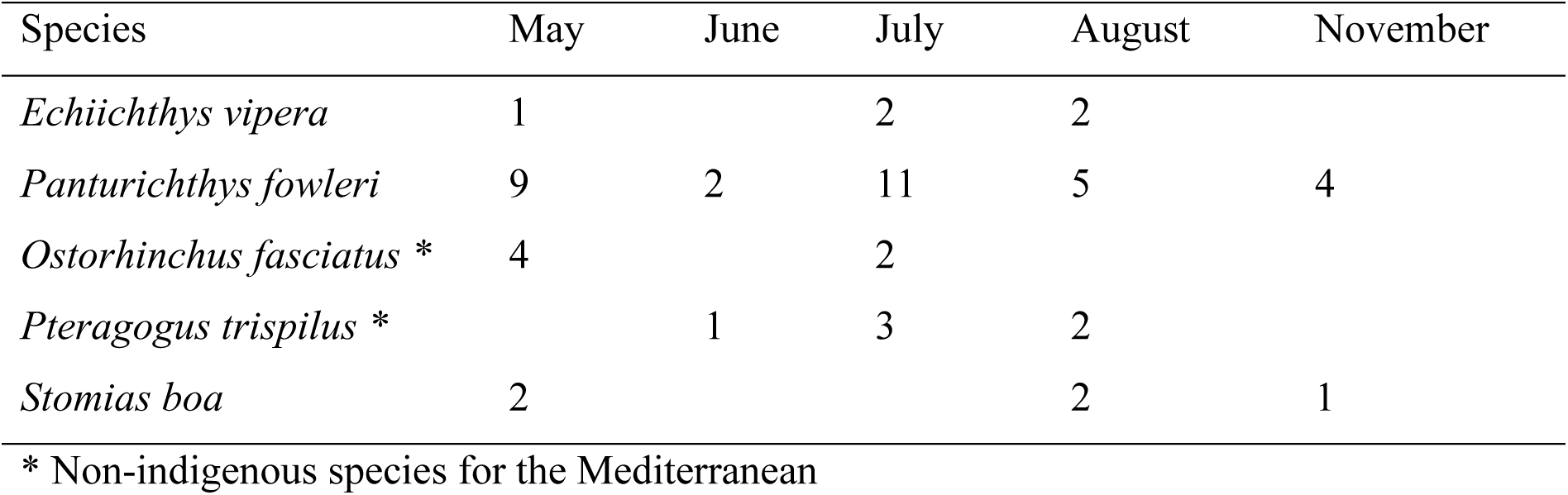
Fish eggs findings of species with no previous knowledge on spawning seasonality.

| Species | May | June | July | August | November |
| --- | --- | --- | --- | --- | --- |
| *Echiichthys vipera* | 1 |  | 2 | 2 |  |
| *Panturichthys fowleri* | 9 | 2 | 11 | 5 | 4 |
| *Ostorhinchus fasciatus* * | 4 |  | 2 |  |  |
| *Pteragogus trispilus* * |  | 1 | 3 | 2 |  |
| *Stomias boa* | 2 |  |  | 2 | 1 |

\* Non-indigenous species for the Mediterranean

### Discussion (Supplementary section S2)

Due to the profound lack of knowledge of Israeli Mediterranean ichthyoplankton and through this study we managed to provide novel knowledge on the distribution and species communities of coastal fish eggs and larvae. Summarizing our results have led to three major conclusions:

1. The coastal ichthyoplankton distribute relatively homogenously throughout the Israeli coastlines. No species-specific assemblages of ichthyoplankton have been found among the different sites, distances from the coast or water columns.
2. Sampling coastal ichthyoplankton can provide new knowledge on the presence of both invasive, cryptic or deep-sea species.
3. Sampling ichthyoplankton can provide new knowledge on spawning seasonality that can in turn improve our ecological and biological knowledge on Mediterranean fauna and can be later implemented in conservation or sustainable fishing activities.

**Table B4.**
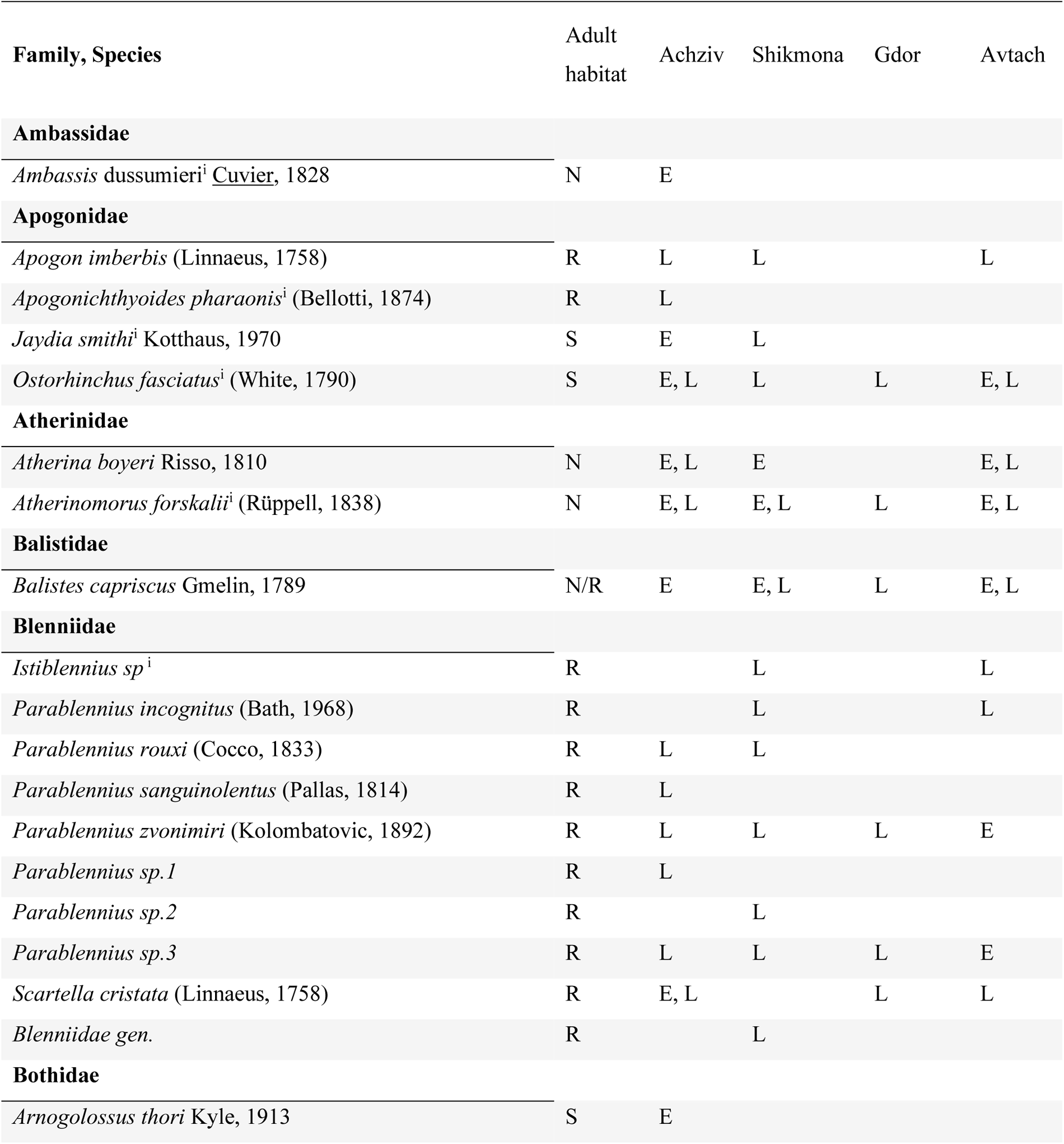
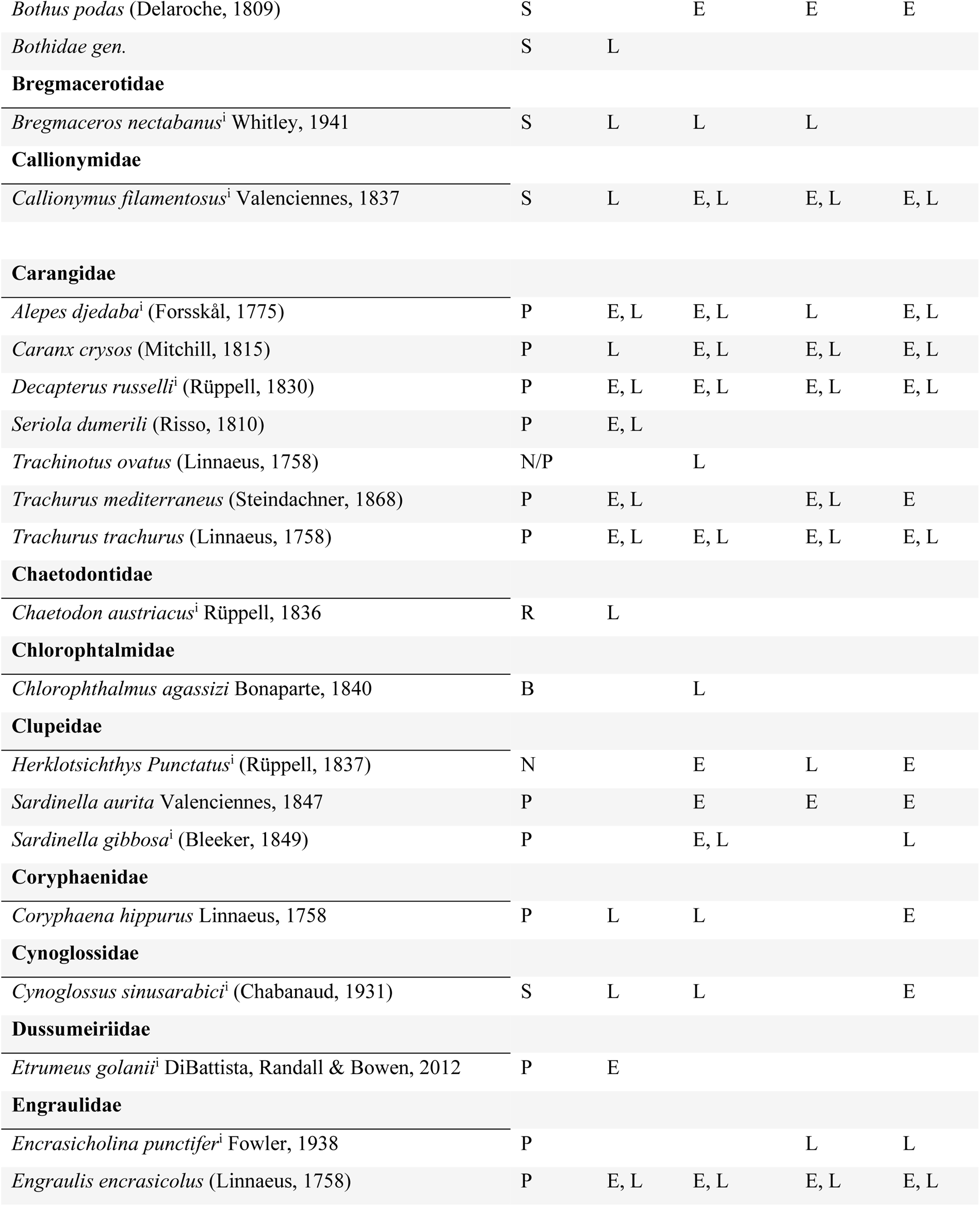
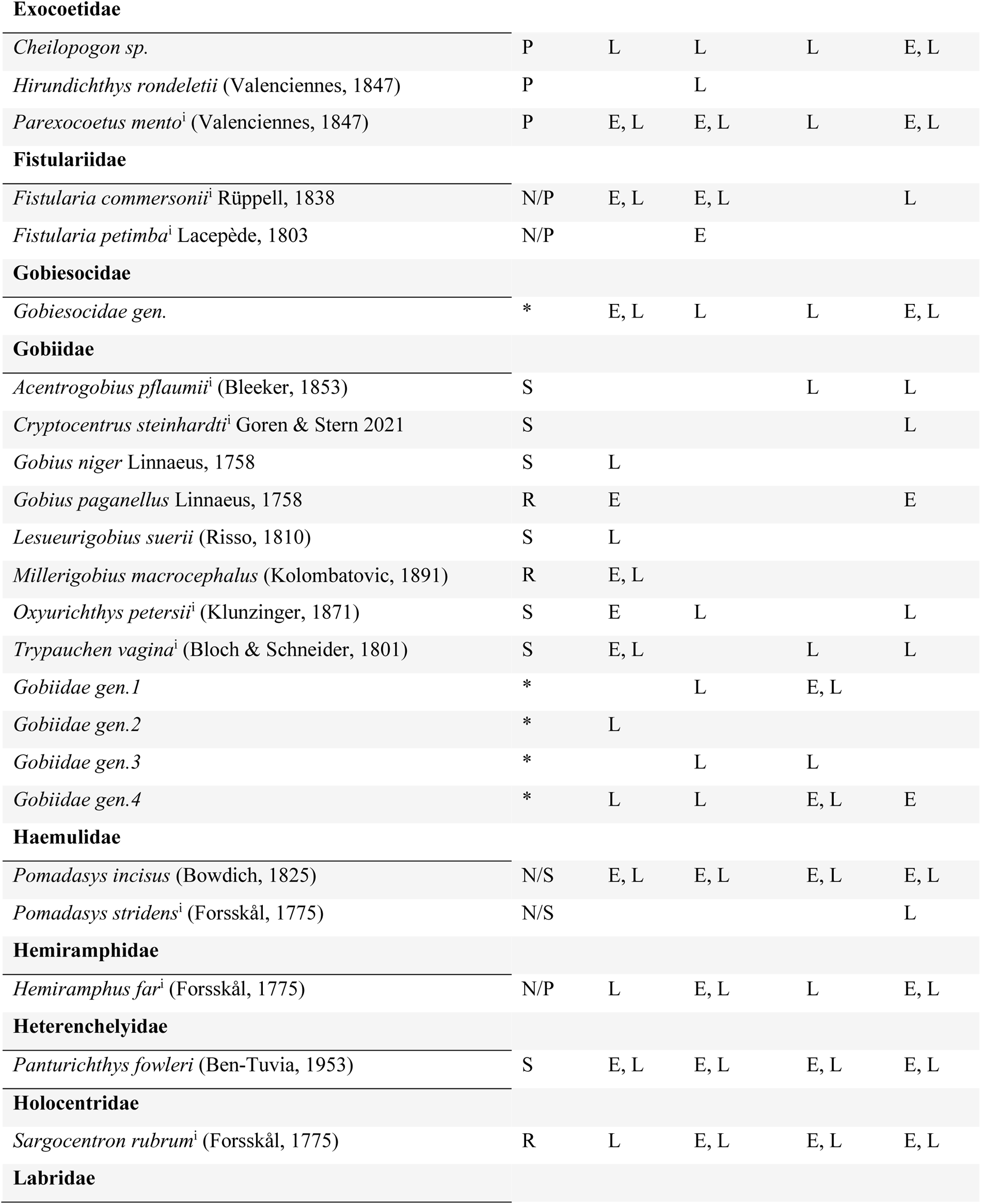
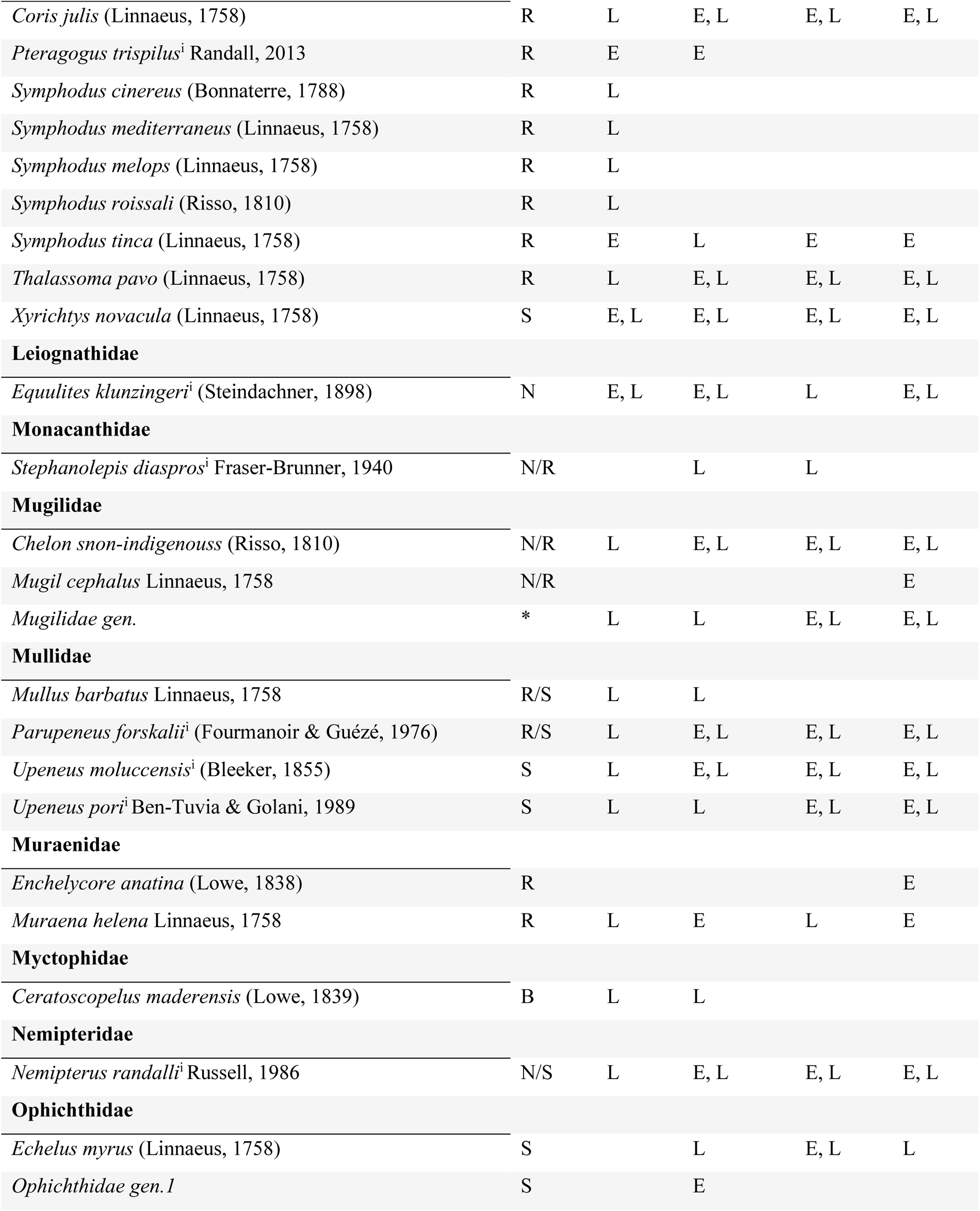
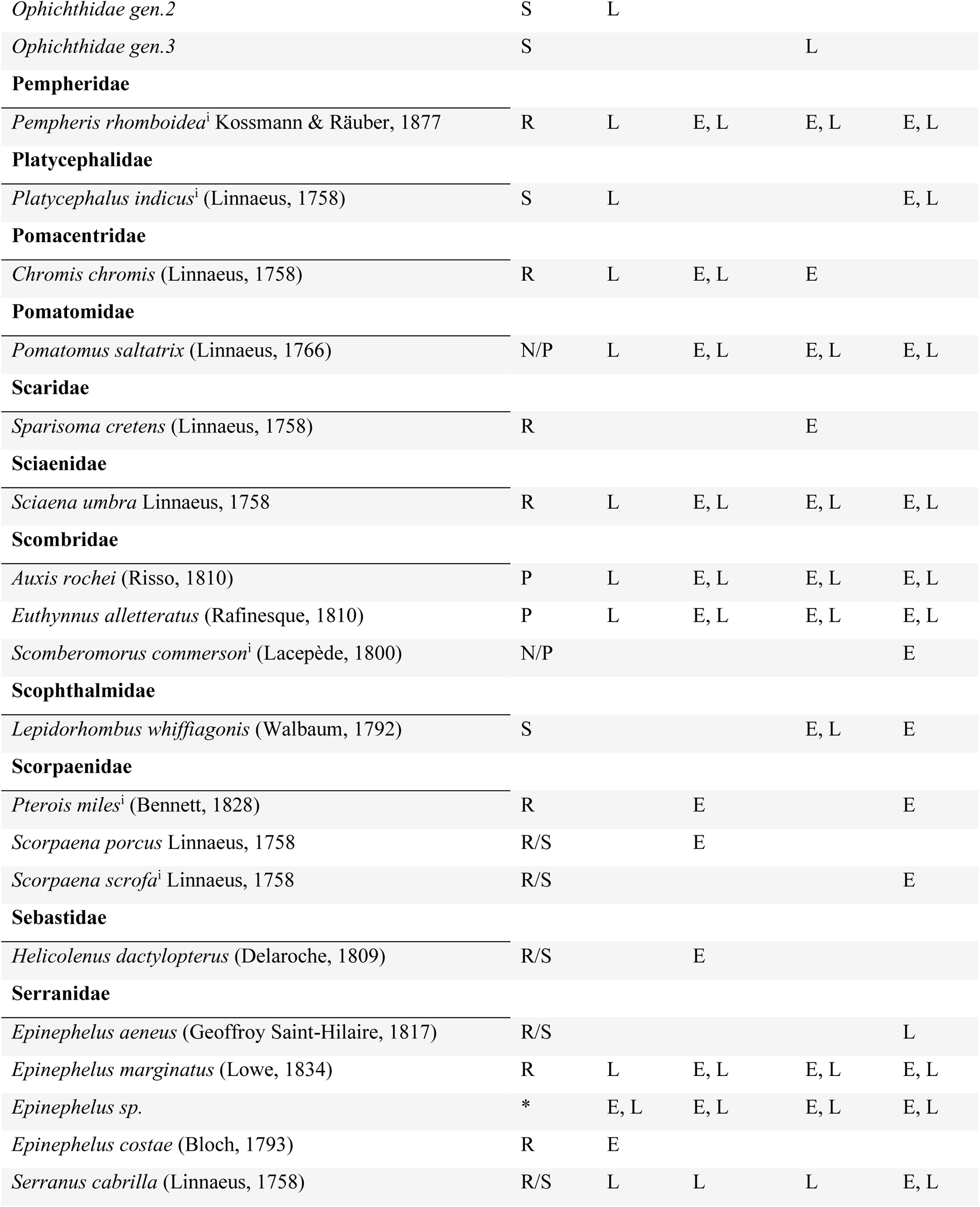
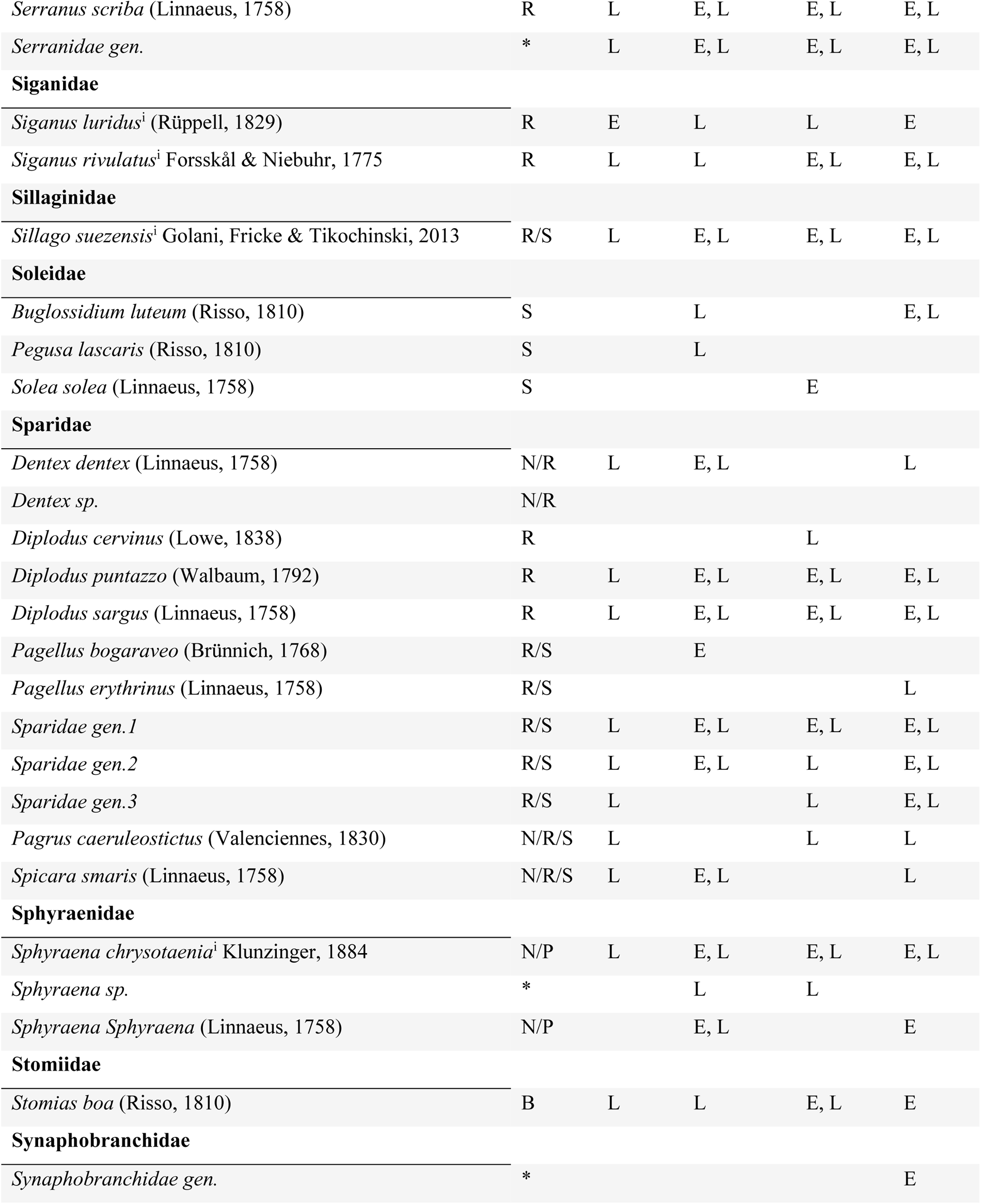
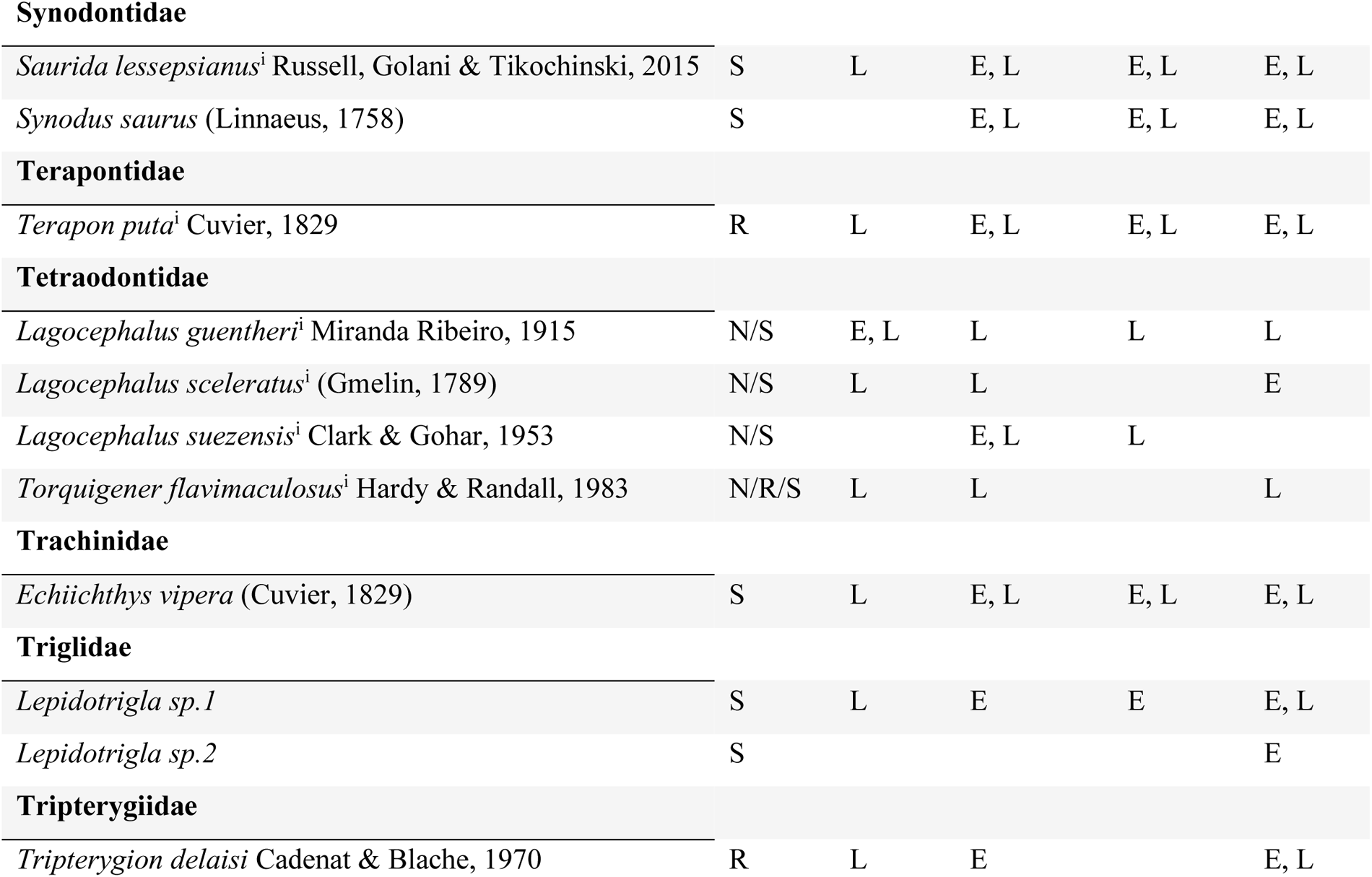
The complete ichthyoplankton inventory found in the different sites of this study, classified to their adult habitat and development stage. B = bathyal habitat; N = neritic habitat; P = pelagic habitat; R = rocky habitat; S = sandy habitat; E = egg stage; L = larval stage. ^1^ = Non-indigenous species

| Family, Species | Adult habitat | Achziv | Shikmona | Gdor | Avtach |
| --- | --- | --- | --- | --- | --- |
| **Ambassidae** |  |  |  |  |  |
| *Ambassis dussumieri*<sup>i</sup> Cuvier, 1828 | N | E |  |  |  |
| **Apogonidae** |  |  |  |  |  |
| *Apogon imberbis* (Linnaeus, 1758) | R | L | L |  | L |
| *Apogonichthyoides pharaonis*<sup>i</sup> (Bellotti, 1874) | R | L |  |  |  |
| *Jaydia smithi*<sup>i</sup> Kotthaus, 1970 | S | E | L |  |  |
| *Ostorhinchus fasciatus*<sup>i</sup> (White, 1790) | S | E, L | L | L | E, L |
| **Atherinidae** |  |  |  |  |  |
| *Atherina boyeri* Risso, 1810 | N | E, L | E |  | E, L |
| *Atherinomorus forskalii*<sup>i</sup> (Rüppell, 1838) | N | E, L | E, L | L | E, L |
| **Balistidae** |  |  |  |  |  |
| *Balistes capriscus* Gmelin, 1789 | N/R | E | E, L | L | E, L |
| **Blenniidae** |  |  |  |  |  |
| *Istiblennius sp*<sup>i</sup> | R |  | L |  | L |
| *Parablennius incognitus* (Bath, 1968) | R |  | L |  | L |
| *Parablennius rouxi* (Cocco, 1833) | R | L | L |  |  |
| *Parablennius sanguinolentus* (Pallas, 1814) | R | L |  |  |  |
| *Parablennius zvonimiri* (Kolombatovic, 1892) | R | L | L | L | E |
| *Parablennius sp.1* | R | L |  |  |  |
| *Parablennius sp.2* | R |  | L |  |  |
| *Parablennius sp.3* | R | L | L | L | E |
| *Scartella cristata* (Linnaeus, 1758) | R | E, L |  | L | L |
| *Blenniidae gen.* | R |  | L |  |  |
| **Bothidae** |  |  |  |  |  |
| *Arnoglossus thori* Kyle, 1913 | S | E |  |  |  |
| *Bothus podas* (Delaroche, 1809) | S |  | E | E | E |
| *Bothidae* gen. | S | L |  |  |  |
| **Bregmacerotidae** |  |  |  |  |  |
| *Bregmaceros nectabanus*<sup>i</sup> Whitley, 1941 | S | L | L | L |  |
| **Callionymidae** |  |  |  |  |  |
| *Callionymus filamentosus*<sup>i</sup> Valenciennes, 1837 | S | L | E, L | E, L | E, L |
| **Carangidae** |  |  |  |  |  |
| *Alepes djedaba*<sup>i</sup> (Forsskål, 1775) | P | E, L | E, L | L | E, L |
| *Caranx crysos* (Mitchill, 1815) | P | L | E, L | E, L | E, L |
| *Decapterus russelli*<sup>i</sup> (Rüppell, 1830) | P | E, L | E, L | E, L | E, L |
| *Seriola dumerili* (Risso, 1810) | P | E, L |  |  |  |
| *Trachinotus ovatus* (Linnaeus, 1758) | N/P |  | L |  |  |
| *Trachurus mediterraneus* (Steindachner, 1868) | P | E, L |  | E, L | E |
| *Trachurus trachurus* (Linnaeus, 1758) | P | E, L | E, L | E, L | E, L |
| **Chaetodontidae** |  |  |  |  |  |
| *Chaetodon austriacus*<sup>i</sup> Rüppell, 1836 | R | L |  |  |  |
| **Chlorophthalmidae** |  |  |  |  |  |
| *Chlorophthalmus agassizi* Bonaparte, 1840 | B |  | L |  |  |
| **Clupeidae** |  |  |  |  |  |
| *Herklotsichthys punctatus*<sup>i</sup> (Rüppell, 1837) | N |  | E | L | E |
| *Sardinella aurita* Valenciennes, 1847 | P |  | E | E | E |
| *Sardinella gibbosa*<sup>i</sup> (Bleeker, 1849) | P |  | E, L |  | L |
| **Coryphaenidae** |  |  |  |  |  |
| *Coryphaena hippurus* Linnaeus, 1758 | P | L | L |  | E |
| **Cynoglossidae** |  |  |  |  |  |
| *Cynoglossus sinusarabici*<sup>i</sup> (Chabanaud, 1931) | S | L | L |  | E |
| **Dussumeiriidae** |  |  |  |  |  |
| *Etrumeus golanii*<sup>i</sup> DiBattista, Randall & Bowen, 2012 | P | E |  |  |  |
| **Engraulidae** |  |  |  |  |  |
| *Encrasicholina punctifer*<sup>i</sup> Fowler, 1938 | P |  |  | L | L |
| *Engraulis encrasicolus* (Linnaeus, 1758) | P | E, L | E, L | E, L | E, L |
| **Exocoetidae** |  |  |  |  |  |
| *Cheilopogon sp.* | P | L | L | L | E, L |
| *Hirundichthys rondeletii* (Valenciennes, 1847) | P |  | L |  |  |
| *Parexocoetus mento*<sup>i</sup> (Valenciennes, 1847) | P | E, L | E, L | L | E, L |
| **Fistulariidae** |  |  |  |  |  |
| *Fistularia commersonii*<sup>i</sup> Rüppell, 1838 | N/P | E, L | E, L |  | L |
| *Fistularia petimba*<sup>i</sup> Lacepède, 1803 | N/P |  | E |  |  |
| **Gobiesocidae** |  |  |  |  |  |
| *Gobiesocidae gen.* | * | E, L | L | L | E, L |
| **Gobiidae** |  |  |  |  |  |
| *Acentrogobius pflaumi*<sup>i</sup> (Bleeker, 1853) | S |  |  | L | L |
| *Cryptocentrus steinhardtii*<sup>i</sup> Goren & Stern 2021 | S |  |  |  | L |
| *Gobius niger* Linnaeus, 1758 | S | L |  |  |  |
| *Gobius paganellus* Linnaeus, 1758 | R | E |  |  | E |
| *Lesueurigobius suerii* (Risso, 1810) | S | L |  |  |  |
| *Millerigobius macrocephalus* (Kolombatovic, 1891) | R | E, L |  |  |  |
| *Oxyurichthys petersii*<sup>i</sup> (Klunzinger, 1871) | S | E | L |  | L |
| *Trypauchen vagina*<sup>i</sup> (Bloch & Schneider, 1801) | S | E, L |  | L | L |
| *Gobiidae gen.1* | * |  | L | E, L |  |
| *Gobiidae gen.2* | * | L |  |  |  |
| *Gobiidae gen.3* | * |  | L | L |  |
| *Gobiidae gen.4* | * | L | L | E, L | E |
| **Haemulidae** |  |  |  |  |  |
| *Pomadasys incisus* (Bowdich, 1825) | N/S | E, L | E, L | E, L | E, L |
| *Pomadasys stridens*<sup>i</sup> (Forsskål, 1775) | N/S |  |  |  | L |
| **Hemiramphidae** |  |  |  |  |  |
| *Hemiramphus far*<sup>i</sup> (Forsskål, 1775) | N/P | L | E, L | L | E, L |
| **Heterenchelyidae** |  |  |  |  |  |
| *Panturichthys fowleri* (Ben-Tuvia, 1953) | S | E, L | E, L | E, L | E, L |
| **Holocentridae** |  |  |  |  |  |
| *Sargocentron rubrum*<sup>i</sup> (Forsskål, 1775) | R | L | E, L | E, L | E, L |
| **Labridae** |  |  |  |  |  |
| *Coris julis* (Linnaeus, 1758) | R | L | E, L | E, L | E, L |
| *Pteragogus trispilus*<sup>i</sup> Randall, 2013 | R | E | E |  |  |
| *Symphodus cinereus* (Bonnaterre, 1788) | R | L |  |  |  |
| *Symphodus mediterraneus* (Linnaeus, 1758) | R | L |  |  |  |
| *Symphodus melops* (Linnaeus, 1758) | R | L |  |  |  |
| *Symphodus roissali* (Risso, 1810) | R | L |  |  |  |
| *Symphodus tinca* (Linnaeus, 1758) | R | E | L | E | E |
| *Thalassoma pavo* (Linnaeus, 1758) | R | L | E, L | E, L | E, L |
| *Xyrichtys novacula* (Linnaeus, 1758) | S | E, L | E, L | E, L | E, L |
| **Leiognathidae** |  |  |  |  |  |
| *Equulites klunzingeri*<sup>i</sup> (Steindachner, 1898) | N | E, L | E, L | L | E, L |
| **Monacanthidae** |  |  |  |  |  |
| *Stephanolepis diaspros*<sup>i</sup> Fraser-Brunner, 1940 | N/R |  | L | L |  |
| **Mugilidae** |  |  |  |  |  |
| *Chelon snon-indigenouss* (Risso, 1810) | N/R | L | E, L | E, L | E, L |
| *Mugil cephalus* Linnaeus, 1758 | N/R |  |  |  | E |
| *Mugilidae* gen. | * | L | L | E, L | E, L |
| **Mullidae** |  |  |  |  |  |
| *Mullus barbatus* Linnaeus, 1758 | R/S | L | L |  |  |
| *Parupeneus forskalii*<sup>i</sup> (Fourmanoir & Guézé, 1976) | R/S | L | E, L | E, L | E, L |
| *Upeneus moluccensis*<sup>i</sup> (Bleeker, 1855) | S | L | E, L | E, L | E, L |
| *Upeneus pori*<sup>i</sup> Ben-Tuvia & Golani, 1989 | S | L | L | E, L | E, L |
| **Muraenidae** |  |  |  |  |  |
| *Enchelycore anatina* (Lowe, 1838) | R |  |  |  | E |
| *Muraena helena* Linnaeus, 1758 | R | L | E | L | E |
| **Myctophidae** |  |  |  |  |  |
| *Ceratoscopelus maderensis* (Lowe, 1839) | B | L | L |  |  |
| **Nemipteridae** |  |  |  |  |  |
| *Nemipterus randalli*<sup>i</sup> Russell, 1986 | N/S | L | E, L | E, L | E, L |
| **Ophichthidae** |  |  |  |  |  |
| *Echelus myrus* (Linnaeus, 1758) | S |  | L | E, L | L |
| *Ophichthidae* gen.1 | S |  | E |  |  |
| *Ophichthidae* gen.2 | S | L |  |  |  |
| *Ophichthidae* gen.3 | S |  |  | L |  |
| **Pempheridae** |  |  |  |  |  |
| *Pempheris rhomboidea*<sup>i</sup> Kossmann & Räuber, 1877 | R | L | E, L | E, L | E, L |
| **Platycephalidae** |  |  |  |  |  |
| *Platycephalus indicus*<sup>i</sup> (Linnaeus, 1758) | S | L |  |  | E, L |
| **Pomacentridae** |  |  |  |  |  |
| *Chromis chromis* (Linnaeus, 1758) | R | L | E, L | E |  |
| **Pomatomidae** |  |  |  |  |  |
| *Pomatomus saltatrix* (Linnaeus, 1766) | N/P | L | E, L | E, L | E, L |
| **Scaridae** |  |  |  |  |  |
| *Sparisoma cretens* (Linnaeus, 1758) | R |  |  | E |  |
| **Sciaenidae** |  |  |  |  |  |
| *Sciaena umbra* Linnaeus, 1758 | R | L | E, L | E, L | E, L |
| **Scombridae** |  |  |  |  |  |
| *Auxis rochei* (Risso, 1810) | P | L | E, L | E, L | E, L |
| *Euthynnus alletteratus* (Rafinesque, 1810) | P | L | E, L | E, L | E, L |
| *Scomberomorus commersoni*<sup>i</sup> (Lacepède, 1800) | N/P |  |  |  | E |
| **Scophthalmidae** |  |  |  |  |  |
| *Lepidorhombus whiffiagonis* (Walbaum, 1792) | S |  |  | E, L | E |
| **Scorpaenidae** |  |  |  |  |  |
| *Pterois miles*<sup>i</sup> (Bennett, 1828) | R |  | E |  | E |
| *Scorpaena porcus* Linnaeus, 1758 | R/S |  | E |  |  |
| *Scorpaena scrofa*<sup>i</sup> Linnaeus, 1758 | R/S |  |  |  | E |
| **Sebastidae** |  |  |  |  |  |
| *Helicolenus dactylopterus* (Delaroche, 1809) | R/S |  | E |  |  |
| **Serranidae** |  |  |  |  |  |
| *Epinephelus aeneus* (Geoffroy Saint-Hilaire, 1817) | R/S |  |  |  | L |
| *Epinephelus marginatus* (Lowe, 1834) | R | L | E, L | E, L | E, L |
| *Epinephelus* sp. | * | E, L | E, L | E, L | E, L |
| *Epinephelus costae* (Bloch, 1793) | R | E |  |  |  |
| *Serranus cabrilla* (Linnaeus, 1758) | R/S | L | L | L | E, L |
| *Serranus scriba* (Linnaeus, 1758) | R | L | E, L | E, L | E, L |
| *Serranidae* gen. | * | L | E, L | E, L | E, L |
| **Siganidae** |  |  |  |  |  |
| *Siganus luridus*<sup>i</sup> (Rüppell, 1829) | R | E | L | L | E |
| *Siganus rivulatus*<sup>i</sup> Forsskål & Niebuhr, 1775 | R | L | L | E, L | E, L |
| **Sillaginidae** |  |  |  |  |  |
| *Sillago suezensis*<sup>i</sup> Golani, Fricke & Tikochinski, 2013 | R/S | L | E, L | E, L | E, L |
| **Soleidae** |  |  |  |  |  |
| *Buglossidium luteum* (Risso, 1810) | S |  | L |  | E, L |
| *Pegusa lascaris* (Risso, 1810) | S |  | L |  |  |
| *Solea solea* (Linnaeus, 1758) | S |  |  | E |  |
| **Sparidae** |  |  |  |  |  |
| *Dentex dentex* (Linnaeus, 1758) | N/R | L | E, L |  | L |
| *Dentex* sp. | N/R |  |  |  |  |
| *Diplodus cervinus* (Lowe, 1838) | R |  |  | L |  |
| *Diplodus puntazzo* (Walbaum, 1792) | R | L | E, L | E, L | E, L |
| *Diplodus sargus* (Linnaeus, 1758) | R | L | E, L | E, L | E, L |
| *Pagellus bogaraveo* (Brünnich, 1768) | R/S |  | E |  |  |
| *Pagellus erythrinus* (Linnaeus, 1758) | R/S |  |  |  | L |
| *Sparidae* gen.1 | R/S | L | E, L | E, L | E, L |
| *Sparidae* gen.2 | R/S | L | E, L | L | E, L |
| *Sparidae* gen.3 | R/S | L |  | L | E, L |
| *Pagrus caeruleostictus* (Valenciennes, 1830) | N/R/S | L |  | L | L |
| *Spicara smaris* (Linnaeus, 1758) | N/R/S | L | E, L |  | L |
| **Sphyraenidae** |  |  |  |  |  |
| *Sphyraena chrysotaenia*<sup>i</sup> Klunzinger, 1884 | N/P | L | E, L | E, L | E, L |
| *Sphyraena* sp. | * |  | L | L |  |
| *Sphyraena Sphyraena* (Linnaeus, 1758) | N/P |  | E, L |  | E |
| **Stomiidae** |  |  |  |  |  |
| *Stomias boa* (Risso, 1810) | B | L | L | E, L | E |
| **Synphobranchidae** |  |  |  |  |  |
| *Synphobranchidae* gen. | * |  |  |  | E |
| **Synodontidae** |  |  |  |  |  |
| *Saurida lessepsianus*<sup>i</sup> Russell, Golani & Tikochinski, 2015 | S | L | E, L | E, L | E, L |
| *Synodus saurus* (Linnaeus, 1758) | S |  | E, L | E, L | E, L |
| **Terapontidae** |  |  |  |  |  |
| *Terapon puta*<sup>i</sup> Cuvier, 1829 | R | L | E, L | E, L | E, L |
| **Tetraodontidae** |  |  |  |  |  |
| *Lagocephalus guentheri*<sup>i</sup> Miranda Ribeiro, 1915 | N/S | E, L | L | L | L |
| *Lagocephalus sceleratus*<sup>i</sup> (Gmelin, 1789) | N/S | L | L |  | E |
| *Lagocephalus suezensis*<sup>i</sup> Clark & Gohar, 1953 | N/S |  | E, L | L |  |
| *Torquigener flavimaculosus*<sup>i</sup> Hardy & Randall, 1983 | N/R/S | L | L |  | L |
| **Trachinidae** |  |  |  |  |  |
| *Echiichthys vipera* (Cuvier, 1829) | S | L | E, L | E, L | E, L |
| **Triglidae** |  |  |  |  |  |
| *Lepidotrigla sp.1* | S | L | E | E | E, L |
| *Lepidotrigla sp.2* | S |  |  |  | E |
| **Tripterygiidae** |  |  |  |  |  |
| *Tripterygion delaisi* Cadenat & Blache, 1970 | R | L | E |  | E, L |

### Supplementary section S3: A detailed account of larval connectivity analysis

**Table C1.** Simulations and their released number of virtual larvae. The name code for the sensitivity analysis scenarios is based on the simulated larval traits of Pleagic Larval Duration (PLD), Mortality coefficient (M), Kappa orientation parameter towards the east (K), Terminal Swimming Speeds (S). For example, 20_0_5_30 scenario is characterized by the following parameters: PLD=20 d, M=0, K=5, S=30 cm/s. For species specific traits see Table 1. For species that are associated with Rocky habitat (i.e., *E. costae* and *S. latus*), virtual larvae were released from all polygons, but more (x3 and x2, respectively) larvae were released from Rocky habitats for these two species.

| Scenario | Type | Effective MPAs<br>(x7 reproductive<br>output) | Number of<br>released virtual<br>larvae |
| --- | --- | --- | --- |
| *Nemipterus randalli* | Species-specific | No | 913,250 |
| *Parupeneus forsskali* | Species-specific | No | 808,700 |
| *Mugil cephalus* | Species-specific | No | 400,030 |
| *Epinephelus costae* | Species-specific | No | 496,800 |
| *Scyllarides latus* | Species-specific | No | 56,496 |
| *Nemipterus randalli* | Species-specific | Yes | 2,071,000 |
| *Parupeneus forsskali* | Species-specific | Yes | 1,966,500 |
| *Mugil cephalus* | Species-specific | Yes | 878,690 |
| *Epinephelus costae* | Species-specific | Yes | 1,958,400 |
| *Scyllarides latus* | Species-specific | Yes | 152,980 |
| 2_0_0_0 | Sensitivity Analysis | No | 913,250 |
| 20_0_0_0 | Sensitivity Analysis | No | 913,250 |
| 20_0_2.5_30 | Sensitivity Analysis | No | 913,250 |
| 20_0_5_15 | Sensitivity Analysis | No | 913,250 |
| 20_0_5_30 | Sensitivity Analysis | No | 913,250 |
| 20_0.2_0_0 | Sensitivity Analysis | No | 913,250 |
| 20_0.4_0_0 | Sensitivity Analysis | No | 913,250 |
| 40_0_0_0 | Sensitivity Analysis | No | 913,250 |

